# Prolonged cold exposure enhances regeneration potential in Arabidopsis

**DOI:** 10.64898/2026.04.20.719581

**Authors:** Fu-Yu Hung, Yetkin Caka Ince, Ayako Kawamura, Arika Takebayashi, Yu Chen, Takuya T. Nagae, Akira Iwase, Noriko Takeda-Kamiya, Kiminori Toyooka, Dongbo Shi, Miguel Moreno-Risueno, Keqiang Wu, Keiko Sugimoto

**Affiliations:** RIKEN Center for Sustainable Resource Science; Yokohama, Kanagawa, Japan; Graduated Institute of Biotechnology, National Chung Hsing University; Taichung, Taiwan; Centro de Biotecnología y Genómica de Plantas (Universidad Politécnica de Madrid (UPM) – Instituto Nacional de Investigación y Tecnología Agraria y Alimentaria-CSIC (INIA-CSIC)), Madrid, Spain; Institute of Plant Biology, National Taiwan University; Taipei, Taiwan; Department of Biological Sciences, Graduate School of Science, The University of Tokyo; Tokyo, Japan

**Keywords:** Plant regeneration, Arabidopsis, prolonged cold, epigenetics, histone acetylation

## Abstract

Prolonged cold exposure over winter impacts plant growth and development but its role beyond flowering regulation remains underexplored. In this study, we show that extended cold enhances regenerative capacity, promoting both callus formation and shoot regeneration in Arabidopsis. This enhancement is mediated by the cold-induced AP2/ERF transcription factors C-REPEAT/DRE-BINDING FACTOR 1 (CBF1), CBF2 and CBF3 which interact with the histone acetyltransferase HISTONE ACETYLTRANSFERASE OF THE GNAT FAMILY 1 (HAG1). The CBFs recruit HAG1 to the loci of key regeneration regulators, such as *WUSCHEL-RELATED HOMEOBOX 5* (*WOX5*), to promote their expression *via* histone acetylation. Our findings thus uncover an epigenetic mechanism by which prolonged cold primes plants for enhanced regeneration, highlighting how environmental cues influence developmental plasticity in plants.

## INTRODUCTION

Prolonged winter-like cold exerts profound effects on plant development, enabling seasonal transitions and enhancing plasticity in growth responses (1–3). One of the best-characterized examples is vernalization, in which extended cold exposure promotes flowering through well-defined genetic and epigenetic mechanisms (4–9). Beyond its role in floral induction, prolonged cold has also been implicated in modulating broader aspects of developmental potential. Notably, horticulture practices have long recognized that winter cold enhances regenerative capacity, improving processes such as adventitious root formation and grafting (10–15). Despite these observations, how prolonged cold influences regeneration remains largely unexplored and the molecular mechanisms underlying this seasonal enhancement of regenerative capacity are yet to be elucidated.

During regeneration, somatic cells reprogramme their fate in response to injury, reacquiring pluripotency and giving rise to new organs or repairing damaged tissues (16–24). This regeneration process is orchestrated by a range of transcription factors, such as WOUND INDUCED DEDIFFERENTIATIONs (WINDs), WUSCHEL-RELATED HOMEOBOX 5 (WOX5) and PLETHORAs (PLTs) (16–20, 25–28). Given the pivotal roles of auxin and cytokinin in regeneration, activation of genes encoding their biosynthesis enzymes, such as YUCCAs (YUCs) and LONELY GUY1 (LOG1), also serves as a crucial driver of cellular reprogramming (20, 29–31). In addition to transcriptional regulations, recent studies have highlighted the role of epigenetic mechanisms in controlling plant regeneration (31–36). In particular, histone modifications catalyzed by histone acetyltransferases, such as HISTONE ACETYLTRANSFERASE OF THE GNAT FAMILY 1/GENERAL CONTROL NONDEREPRESSIBLE 5 (HAG1/GCN5) and HAG3, play essential roles in callus formation and shoot regeneration (31–33).

In this study, we established an experimental system to investigate the role of prolonged cold in plant regeneration. Our findings revealed that 6 weeks of cold treatment enhances callus formation and shoot regeneration in Arabidopsis, and this is mediated by HAG1-dependent histone acetylation that permits increased *WOX5* expression in somatic cells.

## RESULTS

### Prolonged cold exposure promotes wound-induced callus formation and shoot regeneration

To investigate how prolonged winter-like cold affects plant regeneration, we simulated winter conditions through prolonged cold treatment. 14-day-old wild-type (WT) Arabidopsis seedlings grown at 22°C were transferred to 4°C for 6 weeks, while a separate group of 14-day-old seedlings grown at 22°C served as control (Figure 1A). Strikingly, seedlings exposed to prolonged cold exhibited enhanced callus formation at wound sites in various organs including petioles, hypocotyls and leaves, compared to control (Figures 1B and 1C). When hypocotyls were excised from roots, they formed adventitious roots and this process was also enhanced by prolonged cold exposure (Figures S1A and S1B). Moreover, when wounded hypocotyls were cultured on cytokinin-rich shoot inducing medium (SIM), cold-treated plants exhibited accelerated shoot regeneration compared to control (Figures 1D, 1E, S1C, and S1D). To assess how the duration of cold exposure impacts regeneration, we subjected seedlings to 4°C for varying periods ranging from 1 day to up to 8 weeks. As shown in Figure S2, we did not observe significant improvement in callus formation or shoot regeneration after up to 3 weeks of cold treatment. In contrast, regeneration capacity increased progressively between 4 and 6 weeks of cold exposure (Figure S2). We did not detect significant differences between 6-week and 8-week-treatments (Figure S2), indicating that the cold-induced enhancement of regeneration reaches a plateau around 6 weeks.

**Figure 1.**
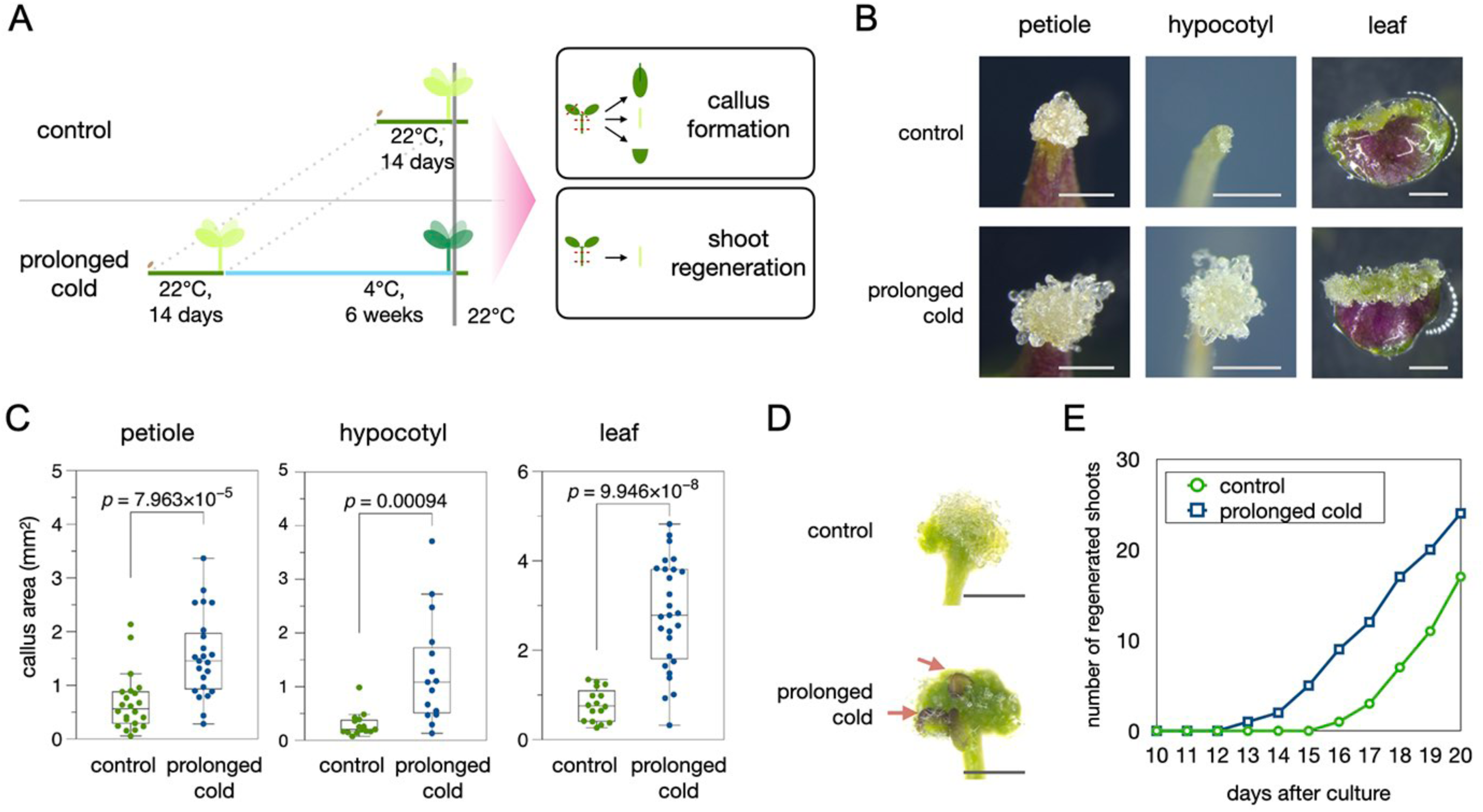
Prolonged cold exposure promotes wound-induced callus formation and shoot regeneration. (A) Experimental workflow of the prolonged cold treatment. 14-day-old seedlings grown at 22°C were transferred to 4°C for 6 weeks and another group of 14-day-old seedlings without cold treatment served as control. (B) Light microscope images of callus forming from wounded wild-type (WT) leaf petioles, hypocotyls and leaf blades. Scale bars, 1 mm. (C) Quantitative data of callus area at 10 days after wounding. The *p*-values were measured by student’s *t*-test. (D) Light microscope images of regenerating shoots from hypocotyl explants cultured on shoot inducting medium (SIM) for 16 days. Red arrows indicated regenerating shoots. Scale bars, 1 mm. (E) Quantitative data of shoot regeneration up to 20 days on SIM.

Previous studies reported that the regenerative capacity varies among *Arabidopsis thaliana* accessions (37). The commonly used Columbia (Col), for instance, exhibits relatively limited regeneration potential whereas Wassilewskija (Ws) displays a stronger ability to regenerate (38, 39). Consistent with these findings, we observed that under standard culture conditions, Ws formed larger callus and regenerated more shoots on SIM compared to Col and Landsberg *erecta* (L*er*) (Figure S3). Notably, all these genotypes showed a significant enhancement in these regenerative traits following 6 weeks of cold treatment (Figure S3), demonstrating that prolonged cold treatment can improve regenerative potential even in genotypes with inherently low regeneration capacity.

### The cold-induced CBF transcription factors associate with HAG1 in plant cells

To investigate the molecular mechanisms underlying the enhancement of plant regeneration by prolonged cold, we first examined the involvement of FRIGIDA (FRI), a central regulator that functions in vernalization-induced flowering (8, 9). *FRI* is highly expressed during the vegetative stage but is repressed after prolonged cold exposure. Notably, the *FRI* transcript in WT Col is truncated and non-functional (9). To assess whether the FRI-mediated pathway contributes to regeneration, we compared WT Col with *FRI* Col, a backcrossed line expressing a functional *FRI* allele (*FRI-sf2*) in the Col background (8). As shown in Figure S4, we found no significant differences in wound-induced callus formation and shoot regeneration, suggesting that the FRI-dependent vernalization pathway is not involved in the prolonged cold-enhanced regenerative response.

To identify key regulators involved in prolonged cold-enhanced regeneration, we performed RNA sequencing (RNA-seq) analysis for the control and prolonged cold treated seedlings. Using this dataset, we examined the expression of 431 regeneration-associated genes we manually compiled based on previous functional and transcriptional studies of regeneration (Table S1) (19). Among these genes, we found that 102 genes were significantly upregulated after 6 weeks of cold treatment based on the criteria of log_2_ fold change > 1 and *p-*value < 0.05 as determined by DEseq2 (Table S1). To predict potential upstream regulators of these prolonged cold-induced, regeneration-associated genes, we applied the gene regulatory network (GRN) inference tool TF2Network (40) and identified 256 candidate transcription factors (Table S2). We further refined this list by selecting transcription factors that have been previously implicated in transcriptional activation and are also upregulated following prolonged cold treatment (Table S2). Among these candidates, members of the *C-REPEAT BINDING FACTORS/DEHYDRATION-RESPONSIVE ELEMENT-BINDING PROTEINS* (*CBF*/*DREB*) family, *CBF1, CBF2* and *CBF3*, caught our attention since the transcript levels of all these *CBF* members were significantly elevated following prolonged cold exposure (Figure 2A). Furthermore, reanalysis of previously published DAP-seq data (41) revealed that the general binding intensity of these CBF transcription factors were strongly enriched at the upstream regulatory regions of prolonged cold-induced regeneration-associated genes (Figures 2B and S5A). Our qPCR analysis further showed that the expression of *CBF1, CBF2* and *CBF3* is negligible under control condition, moderately promoted after 3 weeks of cold exposure and markedly increased after 6 weeks of cold treatment (Figure S5B). These findings suggest that these CBF transcription factors may play a role in mediating the transcriptional upregulation of regeneration genes in response to prolonged cold.

**Figure 2.**
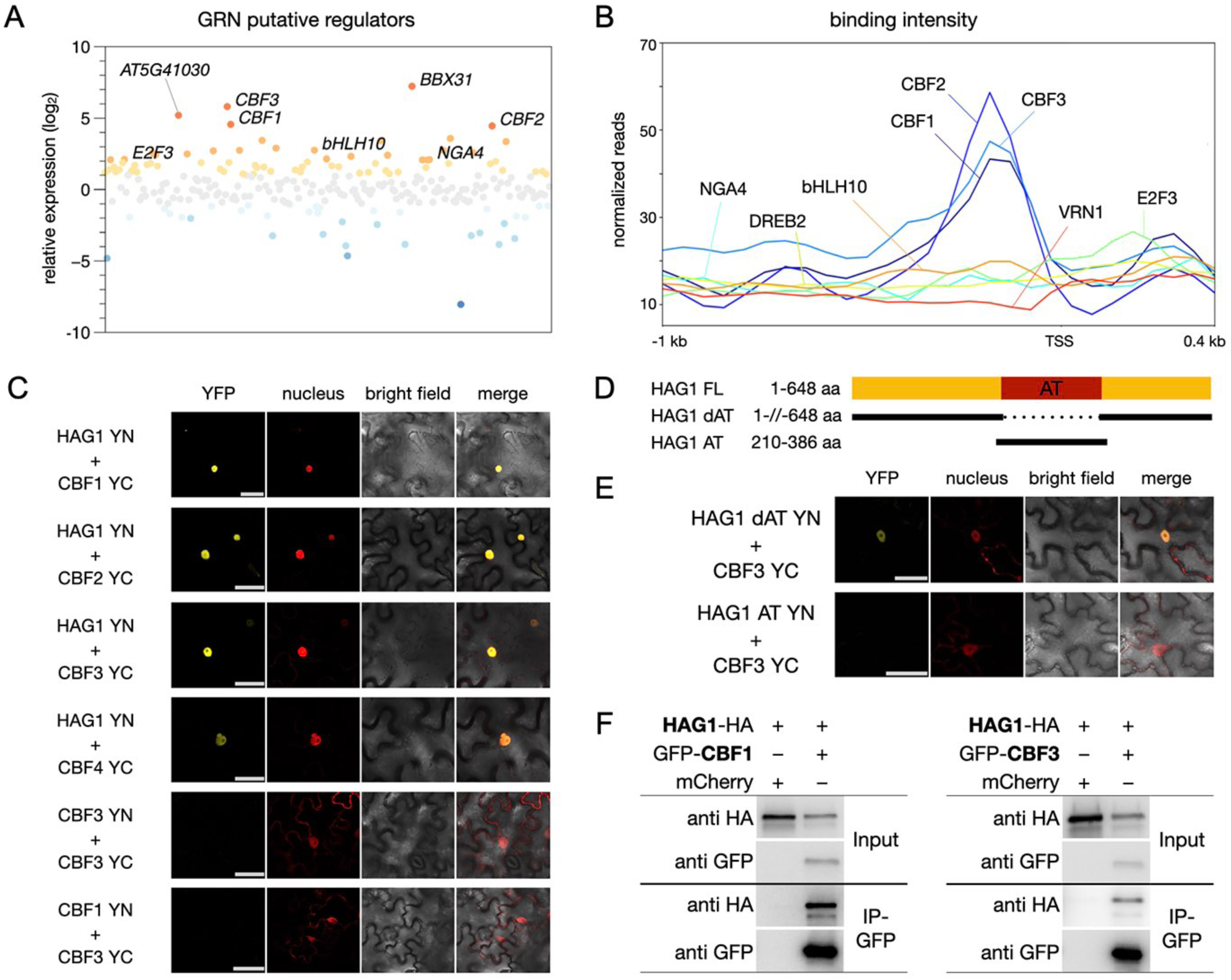
The cold-induced CBF transcription factors associate with HAG1 in plant cells. (A) A scatterplot showing the log2 relative expression level of putative upstream regulators of the cold-enhanced regeneration genes between control and prolonged cold treated plants. (B) A metagene plot showing the overall binding intensity of the selected transcriptional activators on the cold-enhanced regeneration genes. The binding data were extracted from DAP-seq datasets (41). (C) Bimolecular fluorescence complementation (BiFC) assays in *N. benthamiana* leaf cells showing the interaction of HAG1 with CBF1, CBF2 and CBF3. HAG1 and CBF proteins were fused with the N terminus (YN) or C terminus (YC) of YFP and co-delivered into *N. benthamiana* leaves. The nucleus was marked by mCherry carrying a nuclear localization signal. Scale bars, 50 μm. (D) The schematic representation showing the deletions in HAG1 constructs. In the HAG1 dAT constructs, the acetyltransferase (AT) domain was removed from the HAG1 full length (FL) constructs. The HAG1 AT constructs retained only the AT domain. (E) BiFC assays showing that the AT domain is dispensable for the interaction with CBF3. Scale bars, 50 μm. (F) Co-IP experiments of HAG1-HA with CBF1-GFP, CBF3-GFP or mCherry (as control) in transformed *N. benthamiana* leaves. Western blot (WB) experiments were performed with the indicated antibodies.

It is also known that gene expression during plant regeneration is regulated by several histone modification-associated mechanisms (31–36). To examine how these epigenetic marks are affected by prolonged cold exposure, we analyzed the global distribution of histone acetylation (H3Ac), histone H3 lysine 4 trimethylation (H3K4me3), and histone H3 lysine 27 trimethylation (H3K27me3) by chromatin immunoprecipitation followed by sequencing (ChIP-seq) and compared these profiles with gene expression changes following 6 weeks of cold treatment. Previous studies showed that these histone modifications are broadly correlated with transcriptional activation (e.g., H3Ac and H3K4me3) or repression (e.g., H3K27me3) although individual genes are not necessarily regulated by all of these epigenetic marks (36, 42). Consistently, overall levels of H3Ac and H3K4me3 were positively correlated with gene expression changes, while H3K27me3 level was negatively correlated (Figure S6). When we focused on the regeneration-associated genes (Table S1), however, H3Ac exhibited the strongest correlation with expression changes (Figure S6, Table S3), implying that histone acetylation plays a role in mediating prolonged cold-enhanced regeneration.

Interestingly, a previous study by Stockinger et al. (43) reported that the CBF1 transcription factor directly interacts with the histone acetyltransferase HAG1 using *in vitro* pull-down assays (43). Consistently, our *in vivo* bimolecular fluorescence complementation (BiFC) assays, performed in *Nicotiana benthamiana* and Arabidopsis leaves, confirmed that CBF1, as well as CBF2, CBF3, and CBF4, associate with HAG1 in plant cells (Figure 2C). Our data further showed that this *in vivo* association is mediated by regions of HAG1 located outside its acetyltransferase (AT) domain (Figures 2D, 2E, and S7A). These associations were further corroborated by co-immunoprecipitation (co-IP) assays in *Nicotiana benthamiana* and Arabidopsis since we detected the HAG1-HA signals when we immunoprecipitated GFP-CBF1 or GFP-CBF3 proteins with anti-GFP antibodies (Figures 2F and S7B). Together, these results strongly suggest that these CBF transcription factors physically associate with HAG1 and functionally cooperate during the prolonged cold-induced regeneration.

### The CBF-HAG1 module is involved in the prolonged cold-induced regeneration

To test whether CBFs and HAG1 are involved in the prolonged cold-enhanced regeneration, we analyzed wound-induced callus formation in the *cbf1/2/3* triple mutants, *cbf1/2/3-97* and *cbf1/2/3-93* (44), and the *hag1-5* mutant. Without prolonged cold treatment, *hag1-5* mutants exhibited compromised callus formation compared to WT whereas *cbf1/2/3* mutants formed callus at levels comparable to WT (Figures 3A and 3B). Following prolonged cold exposure, WT plants showed a significant increase in wounding-induced callus formation but the *hag1-5* and *cbf1/2/3* mutants did not show a similar enhancement (Figures 3A, 3B, S8A, and S8B).

**Figure 3.**
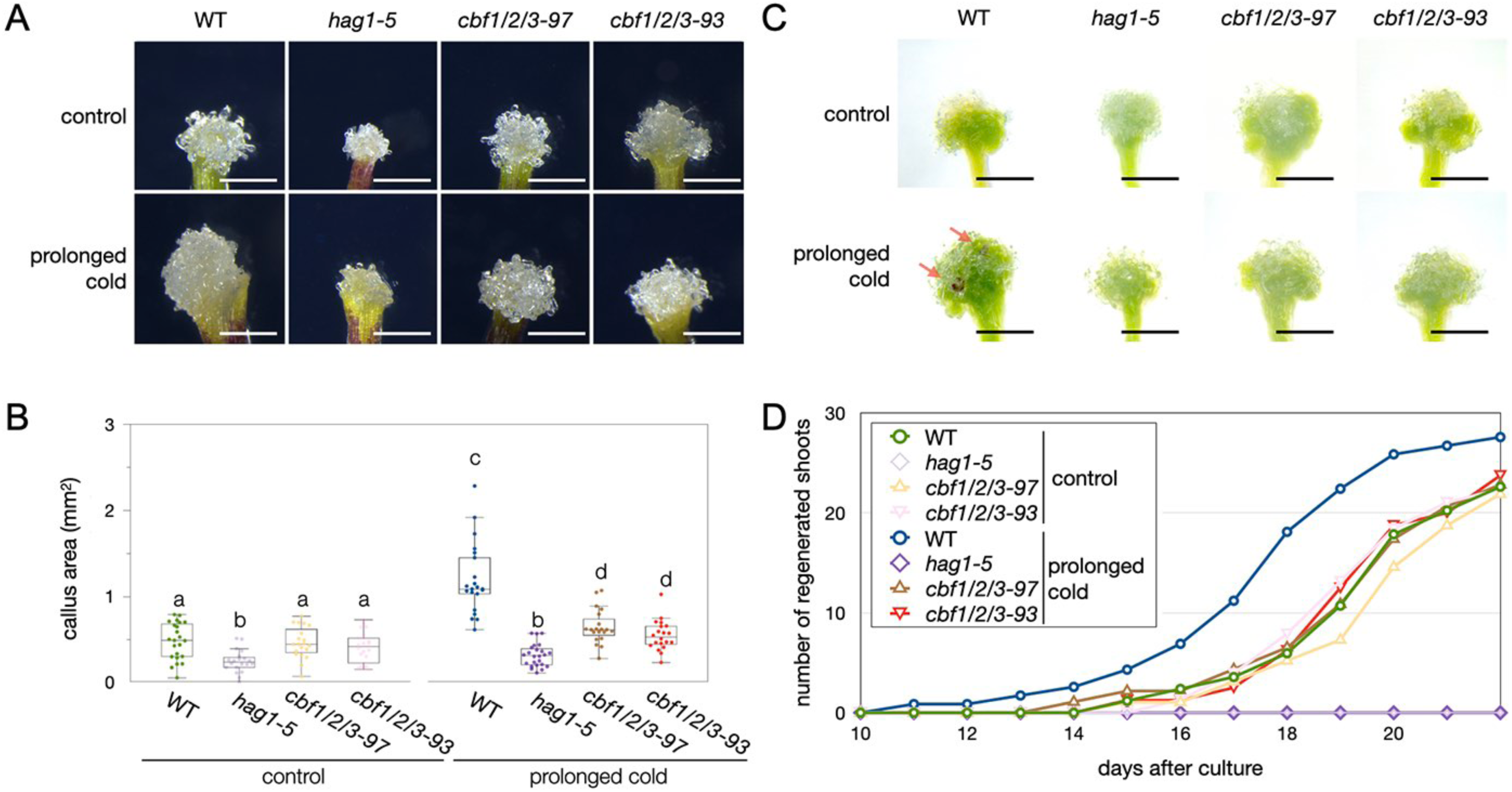
The CBF-HAG1 module is involved in the prolonged cold-induced regeneration. (A-B) Phenotypes (A) and quantitative data (B) of wounding-induced callus in WT, *hag1-5, cbf1/2/3-93,* and *cbf1/2/3-97*. Data were collected at 10 days after wounding. Letters indicated statistical significance based on a two-factor ANOVA with Tukey’s HSD post hoc analysis (*p*□<□0.05). Scale bars, 1 mm. (C, D) Phenotypes (C) and quantitative data (D) of shoot regeneration in WT, *hag1-5, cbf1/2/3-93,* and *cbf1/2/3-97*. Data were collected at 16 days of SIM culture (C, D). Red arrows indicated regenerating shoots. Scale bars, 1 mm.

We further assessed the shoot regeneration in these mutants on SIM. In WT plants, prolonged cold treatment accelerated shoot regeneration by approximately 2 days compared to untreated control. This prolonged cold-induced effect, however, was not observed in *cbf1/2/3* mutants (Figures 3C, 3D, and S8C). The *hag1-5* mutants failed to regenerate shoots under any condition and the prolonged cold treatment did not improve their shoot regeneration capacity (Figures 3C, 3D, and S8C).

Notably, both the callus formation and shoot regeneration phenotypes in *hag1-5* mutants were rescued by expressing HAG1g-GFP in *HAG1pro::HAG1g-GFP hag1-5* lines or HAG1g-FLAG in *HAG1pro::HAG1g-FLAG hag1-5* lines, further confirming that HAG1 is required for prolonged cold-enhanced regeneration (Figures S9A-S9D). In addition, we generated the *hag1 cbf1/2/3* quadruple mutant by crossing *hag1-5* and *cbf1/2/3-97* parental lines. As shown in Figures S9E-S9H, the callus and shoot regeneration phenotypes of the quadruple mutant were comparable to those of *hag1-5* under both control and prolonged cold conditions. Collectively, these results demonstrate that CBF transcription factors and HAG1 function through the same genetic pathway to regulate prolonged cold-enhanced regeneration.

### Prolonged cold-induced gene expression is associated with wounding and cytokinin-mediated pathways

By further characterizing our RNA-seq dataset comparing control and prolonged cold-treated seedlings, we found that prolonged cold treatment led to substantial modification of gene expression with 4607 genes upregulated and 2846 genes downregulated (Table S4). Interestingly, genes typically upregulated by wounding exhibited increased expression following prolonged cold treatment whereas genes downregulated by wounding showed decreased expression (Figure S10), suggesting that prolonged cold amplifies wound-induced transcriptional responses. Since the cold-treated plants were 6 weeks older than control plants, we also examined whether age-related transcriptional changes account for the observed effects. Using the gene sets associated with aging from a previously published study (Table S5, 45), we found no consistent changes in expression that would suggest a significant contribution of tissue aging to the prolonged cold-enhanced regenerative response (Figures S11A and S11B). Morphological analysis also showed no significant differences in aerial tissues between control and prolonged cold-treated seedlings (Figure S11C). These findings strongly suggest that the improved regeneration is unlikely driven by aging-related gene expression, further supporting a direct role for the prolonged cold-induced transcriptional changes in regeneration.

In addition to wound-induced signals, we next assessed whether prolonged cold treatment alters the expression of core cell cycle regulators and meristem-associated genes. We assembled gene sets implicated in cell cycle regulation (46) and meristem development (47) (Table S5) based on previous studies (46, 47). Notably, neither gene group exhibited pronounced global expression changes between control and cold-treated samples (Figure S11D). Together, these findings suggest that prolonged cold treatment alone does not directly activate cell proliferation or meristem formation at the transcriptional level.

Furthermore, plant hormones such as auxin and cytokinin also play key roles in regeneration (10, 16, 18,19, 28–31). To evaluate whether auxin-mediated transcriptional changes contribute to the prolonged-cold promoted regeneration, we compiled a list of auxin- and cytokinin-responsive genes based on previously reported transcriptional datasets (48, 49) (Figures S12A and S12B) (Table S5). The overall expression patterns of these auxin responsive genes, however, did not differ significantly between control and prolonged cold-treated samples (Figure S12A). Consistently, the *auxin response factors 7* (*arf7*) *arf19* double mutants did not exhibit obvious defects in regeneration following prolonged cold exposure (Figures S12C and S12D), suggesting that canonical ARF-mediated auxin signaling is not a major contributor to prolonged cold-induced regeneration.

Similarly, the overall expression of cytokinin-related genes did not show significant changes between control and cold-treated seedlings (Figure S12A). One of the notable exceptions was several *LOG* genes which encodes a key enzyme for cytokinin biosynthesis since their expression was significantly upregulated after 6 weeks of cold treatment (Figure S12B). Furthermore, the *log1/2/3/4/5/7 s*extuple mutants showed significantly reduced regeneration compared to WT after prolonged cold (Figures S12E and S12F), suggesting that the prolonged cold-promoted regeneration is associated with cytokinin-mediated pathways.

### The CBF-HAG1 module regulates the prolonged cold-induced gene expression *via* histone acetylation

To investigate how CBFs and HAG1 regulate the prolonged cold-enhanced regeneration, we additionally performed RNA-seq on *hag1-5* and *cbf1/2/3-97* plants following 6 weeks of cold exposure. By comparing expression profiles between WT, *hag1-5*, and *cbf1/2/3-97* mutants, we found that the prolonged cold-induced transcriptional activation observed in WT was significantly impaired in *hag1-5* and *cbf1/2/3-97* mutants (Figure 4A). We categorized these prolonged cold-induced genes into four groups based on their expression patterns. As shown in Figure 4B, Group 1 consisted of genes that were significantly upregulated in WT but showed diminished induction in both *hag1-5* and *cbf1/2/3-97,* suggesting that their expression depends on both CBFs and HAG1. In contrast, Group 2 and Group 3 included genes that were less induced in either *hag1-5* or *cbf1/2/3-97*, respectively, while Group 4 included genes whose expression was not reduced in either mutant background (Figure 4B). These data indicate that the CBF-HAG1 module plays a key role in activating gene expression following the prolonged cold but additional factors are also involved in this process.

**Figure 4.**
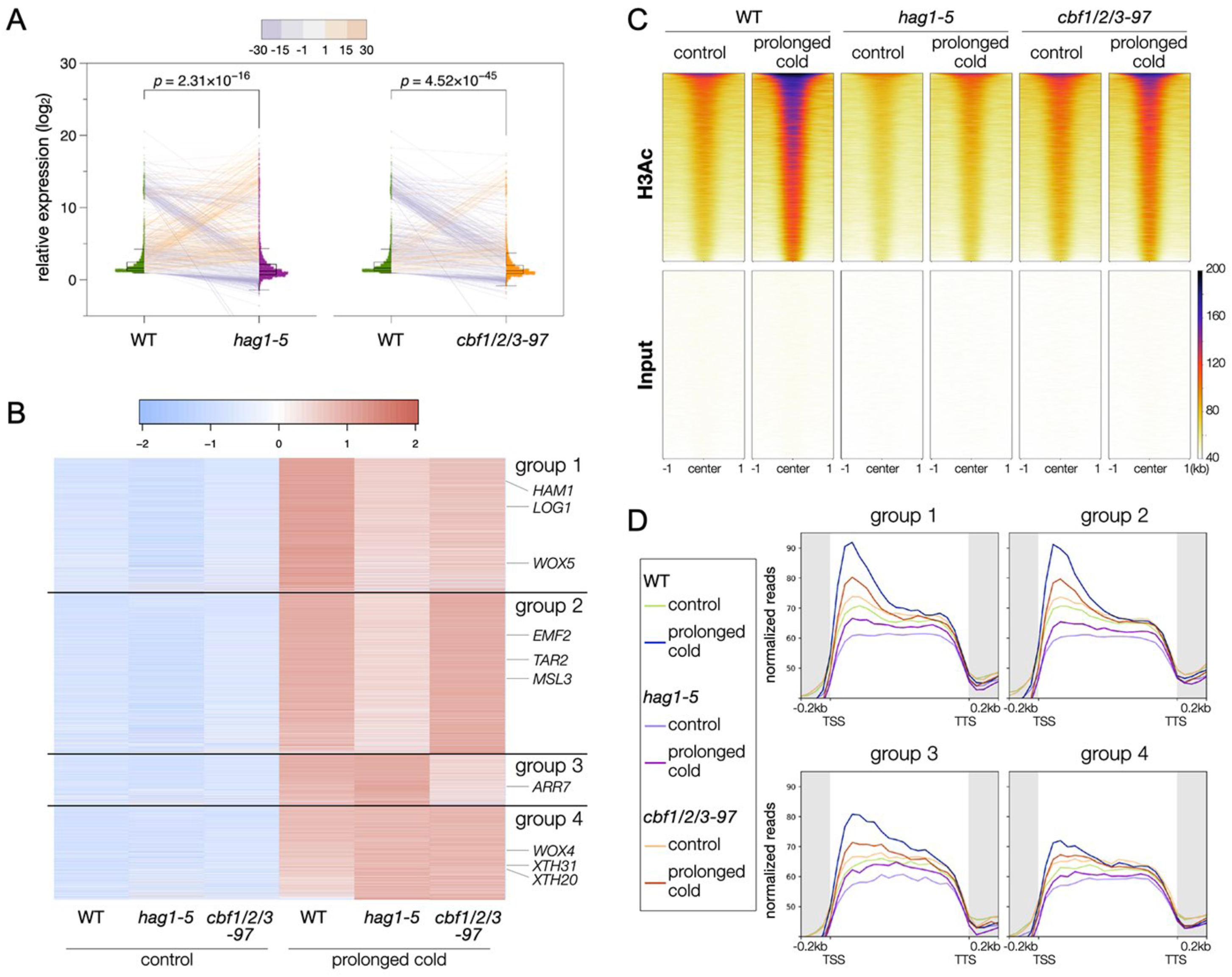
The CBF-HAG1 module regulates the prolonged cold-induced gene expression *via* histone acetylation. (A) The log2 relative expression between control and cold-treated plants in WT, *hag1-5* and *cbf1/2/3-97*. Orange and purple lines indicated increased and decreased, respectively, expression between WT and mutants. (B) Heatmap showing the normalized expression level of the cold-induced genes among WT, *hag1*-5 and *cbf1/2/3-97*. (C) Heatmap showing the relative H3Ac enrichment among WT, *hag1-5* and *cbf1/2/3-97*. The H3Ac levels were presented including 1kb upstream and downstream from the peak center. (D) Metagene profiles showing the H3Ac level of the cold-induced genes among WT, *hag1-5* and *cbf1/2/3-97*. The profile included 0.2□kb upstream of the transcription start site (TSS) to 0.2□kb downstream of the transcription termination site (TTS), and the gene lengths were scaled to the same size.

To determine whether the prolonged cold-induced gene activation is associated with changes in histone acetylation, we subsequently conducted ChIP-seq for H3Ac on WT, *hag1-5*, and *cbf1/2/3-97* plants. Among the genomic regions that exhibited increased H3Ac levels after prolonged cold treatment, we observed a general reduction in H3Ac enrichment in both *hag1-5* and *cbf1/2/3-97* mutants (Figure 4C). Focusing on the 4 gene groups defined earlier (Figure 4B), we found that Group 1 genes exhibited a strong increase in H3Ac levels in WT plants upon the prolonged cold treatment and this increase was markedly diminished in both mutants (Figure 4D). A similar trend was observed for Group 2 genes whose expression also depended on HAG1 (Figure 4D). In contrast, the Group 4 genes, which exhibited HAG1-independent gene activation, showed more modest H3Ac changes in the prolonged cold-treated WT plants (Figure 4D). These results thus show that the CBF-HAG1 module promotes the prolonged cold-induced gene expression in part by modulating H3Ac levels at target loci.

### CBFs recruit HAG1 to the loci of regeneration and developmental regulators to promote their gene expression *via* histone acetylation

Among the genes that showed elevated expression levels associated with increased H3Ac enrichment following prolonged cold exposure, we found several genes previously linked to wound-induced callus formation and/or shoot meristem formation. As shown in Figures 5A and 5B, the expression of these developmental regulators, such as *WOX1*, *WOX5*, *LOG1*, *HAIRY MERISTEM 1* (*HAM1*), *microRNA156c* (*miR156c*) and *PROMOTION OF CELL SURVIVAL 1* (*PCS1*) (19, 50–53), was activated in the prolonged cold-treated WT plants but significantly compromised in the *hag1-5* and *cbf1/2/3-97* mutants, suggesting that these genes are regulated by the CBF-HAG1 module. Consistently, H3Ac enrichment at these loci was also increased in prolonged cold-treated WT, but this increase was not found in the *hag1-5* and *cbf1/2/3-97* mutants (Figure 5C), indicating that this module regulates the expression of these regulators through histone acetylation.

**Figure 5.**
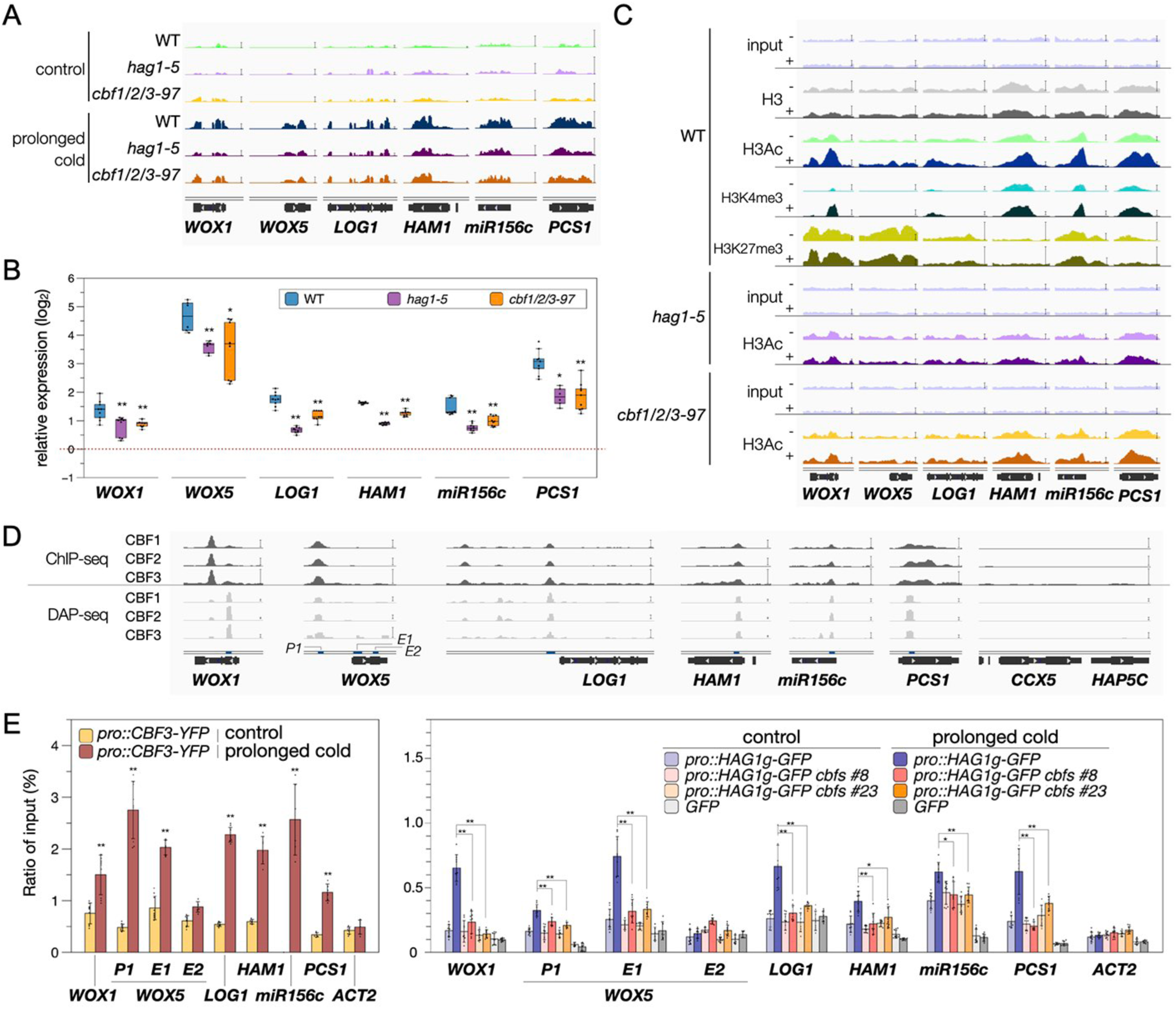
CBFs recruit HAG1 to the loci of several developmental regulators to promote their gene expression *via* histone acetylation. (A) Integrated Genome Viewer showing expression levels of the regeneration-associated genes in WT, *hag1* and *cbf1/2/3-97* under control or after prolonged cold treatment. (B) Relative expression levels of the regeneration-associated genes between control and cold-treated plants in WT, *hag1* and *cbf1/2/3-97*. \**p* < 0.05, \*\**p* < 0.005 (student’s *t*-test). (C) Integrated Genome Viewer showing histone modification levels in WT, *hag1* and *cbf1/2/3-97* under control (-) or after prolonged cold treatment (+). (D) Integrated Genome Viewer showing the binding of CBF1, CBF2 and CBF3 on the regeneration-associated genes. The genomic region of *CCX5* and *HAP5C* was shown as negative control. The binding data were extracted from published ChIP-seq (54) and DAP-seq (41) datasets. The blue lines and letters, *P1*, *E1* and *E2*, indicated the position where primer sets were designed for qPCR in (E). (E) Binding of CBF3-YFP in *CBF3pro::CBF3-YFP* (*pro::CBF3-YFP*) plants (left) and of HAG1-GFP in *HAG1pro::HAG1g-GFP* (*pro::HAG1g-GFP*) and *pro::HAG1g-GFP cbf1/2/3-97* plants (right). ChIP assays were performed with the anti-GFP antibody and the amount of immunoprecipitated DNA was quantified by qPCR. Values represented the normalized average immunoprecipitation efficiencies (%) against the total input DNA. Error bars represent SD. \**p* < 0.05, ** *p* < 0.005 (student’s *t*-test).

These findings support a model in which the CBF-HAG1 module directly drives gene activation in response to prolonged cold. Consistent with this idea, previously published DNA affinity purification sequencing (DAP-seq) (41) and ChIP-seq (54) datasets show that CBF1, CBF2, and CBF3 bind to these target loci both *in vitro* and *in vivo* following short-term cold exposure (Figure 5D). To determine whether CBFs bind to these loci under prolonged cold conditions in our experimental system, we performed chromatin immunoprecipitation followed by quantitative PCR (ChIP-qPCR). Using *CBF3pro::CBF3-YFP* seedlings (55) exposed to 6 weeks of cold, we observed clear enrichment of CBF3-YFP at the target loci compared with control conditions (Figure 5E). ChIP-qPCR analysis using *HAG1pro::HAG1g-GFP* seedlings further revealed that HAG1-GFP also accumulates at these loci after prolonged cold treatment (Figure 5E). Notably, this enhancement was markedly reduced in the *HAG1pro::HAG1g-GFP cbf1/2/3-97* background (Figure 5E). These results together demonstrate that CBFs are required for the recruitment of HAG1 to prolonged cold-responsive loci, thereby linking transcription factor binding with chromatin remodelling to promote regeneration.

### Prolonged cold-induced *WOX5* expression enhances callus formation and subsequent shoot regeneration

Among the developmental regulators induced through histone acetylation, we focused on *WOX5* for further analysis as it showed strong induction at both the transcript and H3Ac levels following prolonged cold treatment (Figure 6A). Under normal conditions *WOX5* expression is confined to the stem cell niche within the root meristem and its expression is repressed in differentiated cells (56). Notably, confocal imaging of *WOX5pro::NLS-GFP* plants (57) revealed ectopic activation of the *WOX5* promoter after 6 weeks of cold treatment, with nuclear GFP signals detected in differentiated hypocotyl cells (Figures 6B and S13A, Video S1). Quantification of GFP-positive nuclei in control, 3-week, and 6-week cold-treated seedlings revealed a slight increase after 3 weeks of cold exposure and a pronounced induction after 6 weeks (Figure S13B). In addition, we observed increased HAG1-GFP and NLS-mCherry signals in the *HAG1pro::HAG1g-GFP* and *CBF3pro::NLS-mCherry* reporter lines, respectively, after 6 weeks of cold treatment (Figure 6C), supporting a model in which elevated accumulation of CBF3 and HAG1 promotes *WOX5* expression. Callus is known to originate from pericycle or vasculature cells in wounded hypocotyls (20). Consistently, the fluorescent signals driven by *HAG1*, *CBF3* or *WOX5* promoters were clearly detected in pericycle cells although they were also present in other cell types of hypocotyls (Figures 6B and 6C).

**Figure 6.**
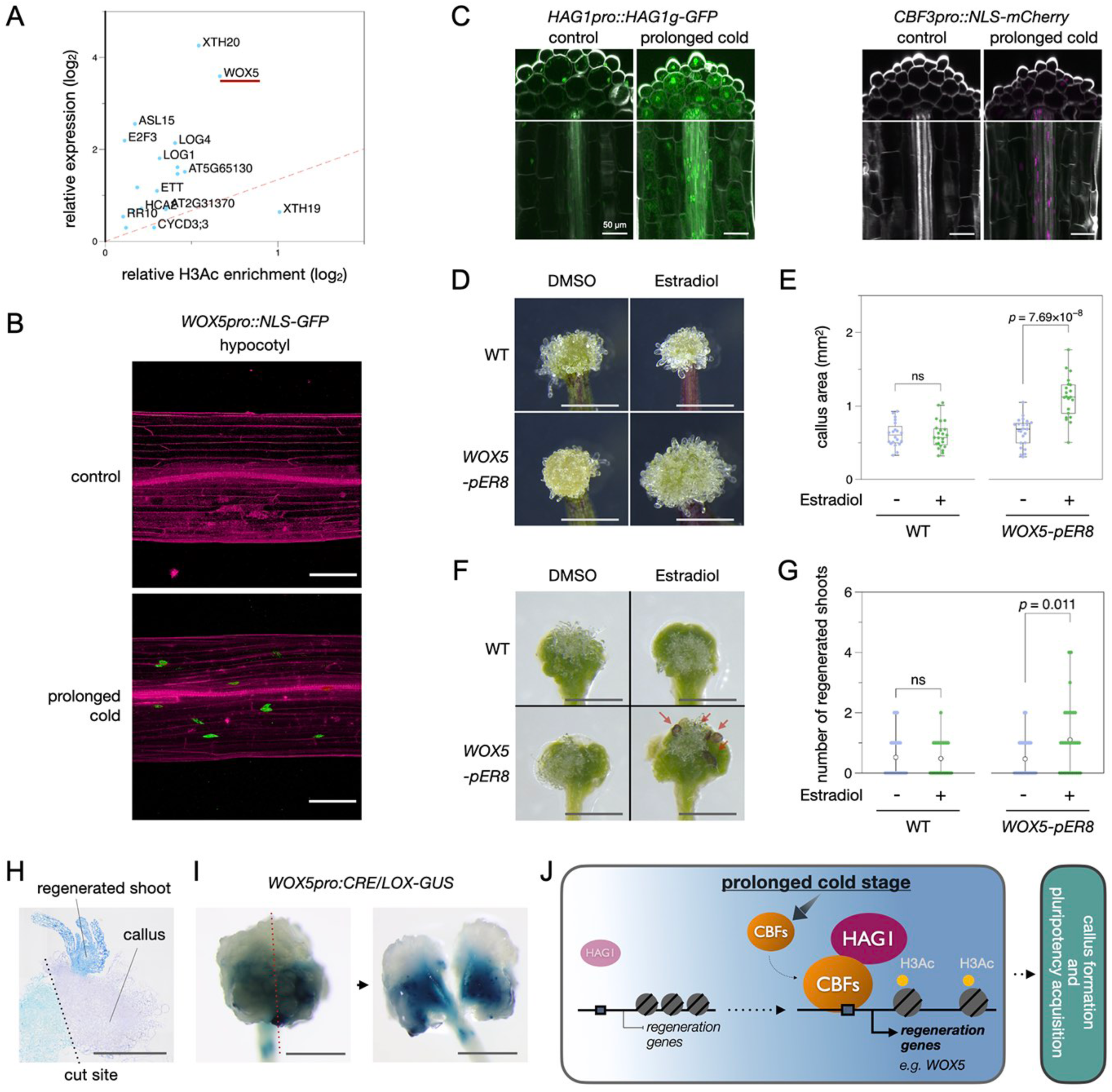
Prolonged cold-induced *WOX5* expression enhances callus formation and subsequent shoot regeneration. (A) X-Y scatter plot showing the regeneration-associated genes activated by prolonged cold treatment with increased H3 acetylation. The abline indicates the correlation between expression levels and H3Ac enrichment. (B) Confocal microscope images of *NLS-GFP* expression driven by the *WOX5* promoter in *WOX5pro::NLS-GFP* reporter lines in control or after prolonged cold treatment. Z-stack images taken from the hypocotyl cells are projected. The GFP signal was marked in green and PI staining was marked in red. Scale bars, 100 μm. (C) Confocal microscope images of *HAG1-GFP* and *NLS-mCherry* expression driven by the *HAG1* and *CBF3* promoters, respectively, in control or after prolonged cold treatment. Scale bars, 50 μm. (D, E) Phenotypes and quantitative data of wounding induced callus in *WOX5-pER8* plants after treatment with DMSO or 10 μM β-estradiol. Scale bars, 1mm. (F, G) Phenotypes and quantitative data of shoot regeneration in *WOX5-pER8* plants after treatment with DMSO or 10 μM β-estradiol. Red arrows indicated regenerating shoots. Scale bars, 1mm. ns, not significant. (I) The light microscope image of a longitudinal section of wound-induced callus after 6 weeks of cold treatment, showing shoot regeneration from callus tissue in a hypocotyl. Scale bar, 1 mm. (J) The light microscope images of callus formed from wounded *WOX5pro:CRE/LOX-GUS* hypocotyls after 6 weeks of cold treatment. Explants were treated with 10 µM DEX after dissection during SIM culture. Callus is bisected to reveal *WOX5* promoter activity within the internal tissue. Scale bar, 1 mm. (K) A schematic diagram showing how prolonged cold promotes callus formation and subsequent shoot regeneration. The cold-induced CBF transcription factors recruit HAG1 to the locus of key regeneration genes, such as *WOX5,* to increase H3 acetylation and consequently gene expression, leading to the increased regeneration potential.

Previous studies reported that increased *WOX5* expression promotes shoot regeneration in Arabidopsis tissue culture (26, 27). Similarly, induced overexpression of *WOX5* enhanced wounding-induced callus formation and subsequent shoot regeneration in *WOX5-pER8* plants treated with estradiol prior to wounding and/or SIM incubation (Figures 6D-G). Conversely, these regenerative responses were compromised in the *wox5-1* mutant compared with WT following prolonged cold treatment (Figures S13C and S13D), further supporting a critical role of WOX5 in prolonged cold-enhanced regeneration. Importantly, we consistently observed that enhanced shoot regeneration was accompanied by increased formation of wound-induced callus. Histological analysis of callus grown on SIM further revealed that shoots regenerated from callus tissues rather than directly from pre-existing hypocotyl tissues (Figure 6H). To determine whether wound-induced callus originates from *WOX5*-expressing cells, we attempted to perform lineage tracing experiments using *WOX5pro:CRE/LOX-GUS* lines in which dexamethasone (DEX)-induced CRE/LOX activity marks cells that have experienced active *WOX5* promoter activity (26, 58). We initially treated these seedlings with 10 µM DEX during 6 weeks of cold treatment to induce transcription of CRE-GR by the *WOX5* promoter. This approach was, however, unsuccessful possibly because CRE recombinase activity is not sufficiently efficient at 4°C to drive the recombination required to convert *35Spro:LOX2272-GFP-ter-LOX2272-GUS-ter* into *35Spro:GUS-ter*. To circumvent this problem, we treated seedlings with DEX after transfer to SIM at 22°C. As shown in Figure 6I, internal tissues within the wound-induced callus exhibited strong and widespread GUS staining, suggesting that callus formation occurs, at least in part, from cells that express *WOX5* gene. Together, the above results support that prolonged cold promotes regeneration by inducing *WOX5* expression, which facilitates wound-induced callus formation and subsequent shoot regeneration.

## DISCUSSION

Our findings demonstrate that extended, winter-like cold exposure enhances regenerative capacities in Arabidopsis. During this process, the CBF transcription factors are upregulated and recruit HAG1 to target loci, resulting in elevated histone H3 acetylation and activation of key regeneration genes, such as *WOX5* (Figure 6J). These prolonged cold-induced epigenetic modifications likely prime plants for enhanced callus formation and shoot regeneration. A recent study by Perez-Garcia et al. (55) revealed that *CBF3* is transcriptionally activated in the root endodermis even under non-cold conditions and CBF3 proteins move into adjacent meristematic tissues including the stem cell niche where it regulates the *WOX5* expression. Our study showed that prolonged cold exposure markedly expands the expression domains of *CBF3* and *HAG1*, accompanied by broader *WOX5* expression in hypocotyl tissues (Figures 6B, 6C, S13). These data further support the functional relationship between CBFs and WOX5 and uncover an additional regulatory layer mediated by HAG1-dependent histone acetylation. The above study also showed that CBF3 is required for *WOX5* activation during root development in response to short-term cold (55), suggesting that CBF-mediated *WOX5* regulation operates across different tissue types. Interestingly, in our system, 3 weeks of cold treatment was insufficient to induce callus formation from hypocotyls (Figure S2), suggesting that additional regulatory barriers restrict regeneration under shorter cold exposure. Our data also highlight the complexity of cold-responsive transcriptional changes, as only a subset of the prolonged cold-enhanced gene expression is mediated by the CBF-HAG1 module (Figure 4B). Further dissection of CBF- or HAG1-independent pathways will be critical to fully elucidate how transcriptional regulators and chromatin modifiers coordinate the response after extended cold treatment.

Previous studies reported that *de novo* organogenesis, such as rooting and *in vitro* regeneration, is more efficient during the winter months across diverse plant species (15, 59–61). Likewise, late winter to early spring is often the most favorable period for grafting, with improved success rates reported in various species (11–13). Since successful grafting is linked to robust callus formation at wound sites (10, 14, 16, 17), these seasonal enhancements in regenerative potential may reflect conserved long-term cold-responsive mechanisms similar to the CBF-HAG1 module identified in our study. Understanding and harnessing these prolonged cold-induced epigenetic pathways could offer potential new strategies to improve regeneration efficiency in plant biotechnology, particularly for species or cultivars that are typically recalcitrant to regeneration.

### Limitations of the study

This study was focused on investigating how prolonged cold exposure primes plant tissue for regeneration, given that plants exposed to prolonged cold do not exhibit overall morphological changes (Figure S11C), suggesting that cold treatment, and by extension the CBF-HAG1 complex, does not directly activate meristem activity or cell proliferation during cold period. This interpretation is supported by our RNA-seq data, which show that expression levels of key meristem regulators, with the exception of *WOX5*, and core cell cycle genes remain largely unchanged right after prolonged cold treatment (Figure S11D). Based on these observations, we propose a model in which the CBF-HAG1 complex primes tissues by enhancing regenerative potential during cold exposure, while injury-derived signals are still required to trigger actual activation of meristem activity and cell proliferation. This is consistent with what is observed in nature where plants do not regenerate overwinter until the proper environmental stimuli were delivered. Future studies should investigate how this primed state influences meristem activity and cell proliferation to facilitate regeneration.

This study was also focused on investigating the role of H3 acetylation but it is also possible that H3K27me3 also plays some roles in cold-enhanced regeneration. While global changes in H3K27me3 do not show a strong correlation with the expression of regeneration-associated genes following prolonged cold exposure (Figure S6), we did identify several genes, including *CBFs*, that exhibit decreased H3K27me3 accompanied by increased expression after 6 weeks of cold treatment (Table S3), implying that H3K27me3 may also contribute to the prolonged cold-mediated regeneration. It is also possible that among the 1047 genes that lose H3K27me3 during 6 weeks of cold, some of them may gain expression at later time points for which we did not have RNA-seq data. Changes in H3K27me3 and/or gene expression might also occur in a small population of cells and these changes were below the detection limit of our bulk RNA-seq and ChIP-seq analyses. Future studies should thus explore the role of H3K27me3 in this regeneration system by using single cell RNA-seq and ChIP-seq as well as inducible loss-of-function mutations in the PRC2 complex.

## RESOURCE AVAILABILITY

### Lead Contact

Further information and requests for resources and reagents should be directed to and will be fulfilled by the Lead Contact, Keiko Sugimoto. The requests for the materials can be also submitted to Fu-Yu Hung.

### Materials Availability

This study did not generate new unique reagents. All strains, seeds and plasmids used in this study are available upon request.

### Data and Code Availability

This paper does not report original code. All data reported in this paper will be shared by the lead contact upon request. The raw data of RNA-seq and ChIP-seq experiments for this study have been deposited in the Gene Expression Omnibus (GEO) database with the accession code ChIPseq: GSE297486 and RNA-seq: GSE297467. Any additional information required to reanalyze the data reported in this paper is available from the lead contact upon request.

## ACKNOWLEDGMENTS

The authors thank Duncan Coleman, Mariko Mouri, Chika Ikeda, Noriko Doi and Mayuko Sato for technical assistance. This study was supported by Ministry of Education, Culture, Sports, Science and Technology grant 20H05911 (K.S.); Japan Society for the Promotion of Science grant 20H03284 (K.S.); Japan Science and Technology Agency GteX grant JPMJGX23B (K.S.); ASPIRE grant JPMJAP2306 (K.S.); Japan Society for the Promotion of Science postdoctoral fellowship (F-Y. H.); and National Science and Technology Council (Taiwan) grant 114-2628-B-005-001 (F-Y. H.). National Science and Technology Council (Taiwan) grants 111-2311-B-002-025-MY3 and 113-2311-B-002-002 (K.W.); and National Taiwan University grants 114L893201 and 114L8521 (K.W.).

## AUTHOR CONTRIBUTIONS

Conceptualization: F-YH and KS. Methodology: F-YH, AT, AK, YCI, AI and YC. Investigation: F-YH, YCI, AK, AT, YC, TN, AI, NT, KT, DS and MM-R. Funding acquisition: F-YH and KS. Supervision: KW and KS. Writing – original draft: F-YH and KS. Writing – review & editing: YCI, AK, AT, YC, TN, AI, NT, KT, DS, MM-R and KW.

## SUPPLEMENTAL INFORMATION

**Document S1. Figures S1-S13**

**Table S1. List of regeneration-associated genes. Related to Figure 2**.

**Table S2. List of putative regulators for prolonged cold-induced expression of regeneration-associated genes. Related to Figures 2 and S5.**

**Table S3. List of loci that change H3ac, H3K4me3 and H3K27me3 levels after prolonged cold exposure. Related to Figure S6.**

**Table S4. List of gene expression patterns in WT, *hag1-5* and *cbf1/2/3-97* following prolonged cold. Related to Figures 4, S10, and S11.**

**Table S5. List of wounding, aging, auxin, CK, cell-cycle, meristem development and flowering-associated genes. Related to Figures S6 and S11.**

**Video S1. *WOX5pro::NLS-GFP* expression. Related to Figures 6 and S13.**

## STAR⍰METHODS

### KEY RESOURCES TABLE

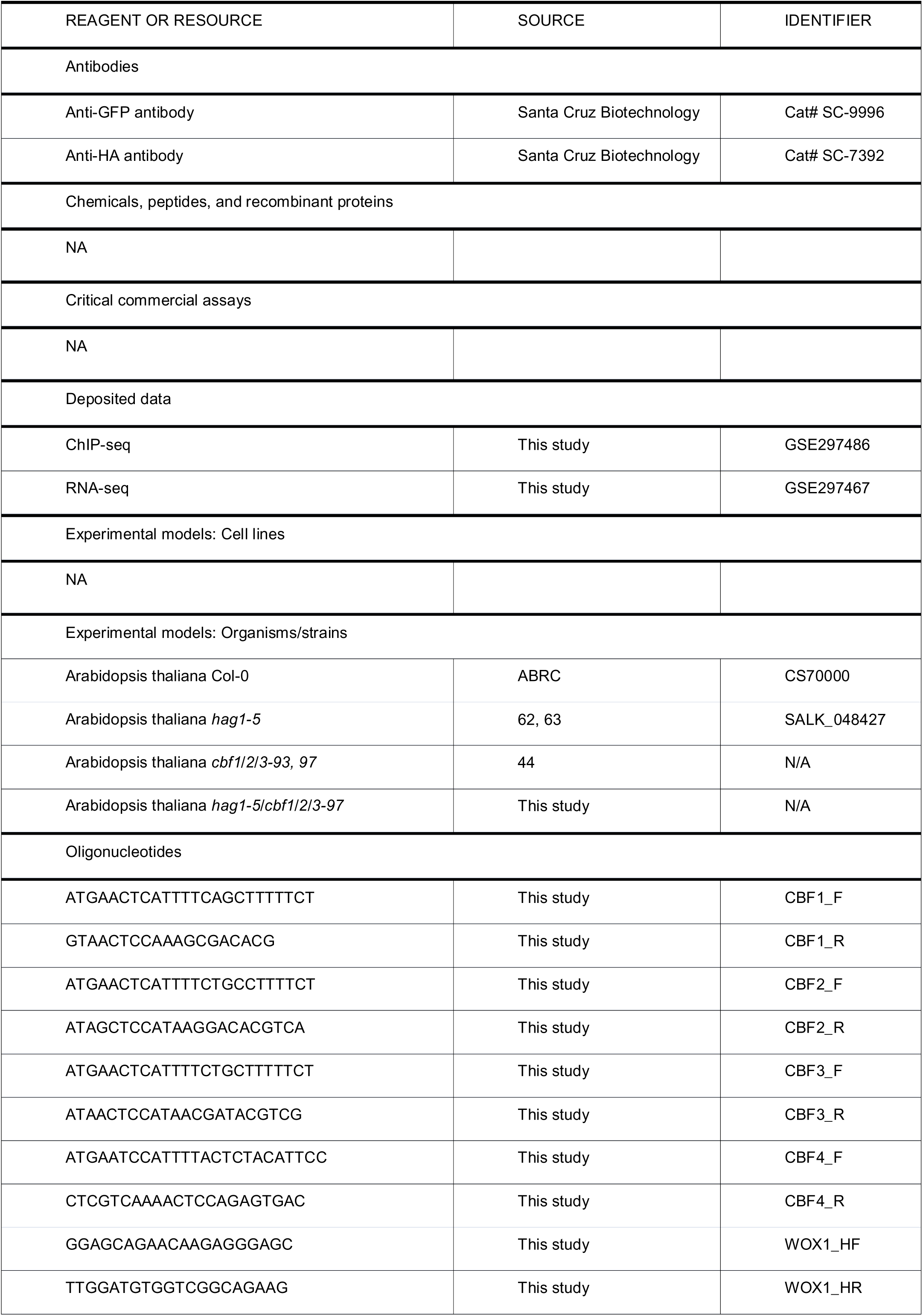
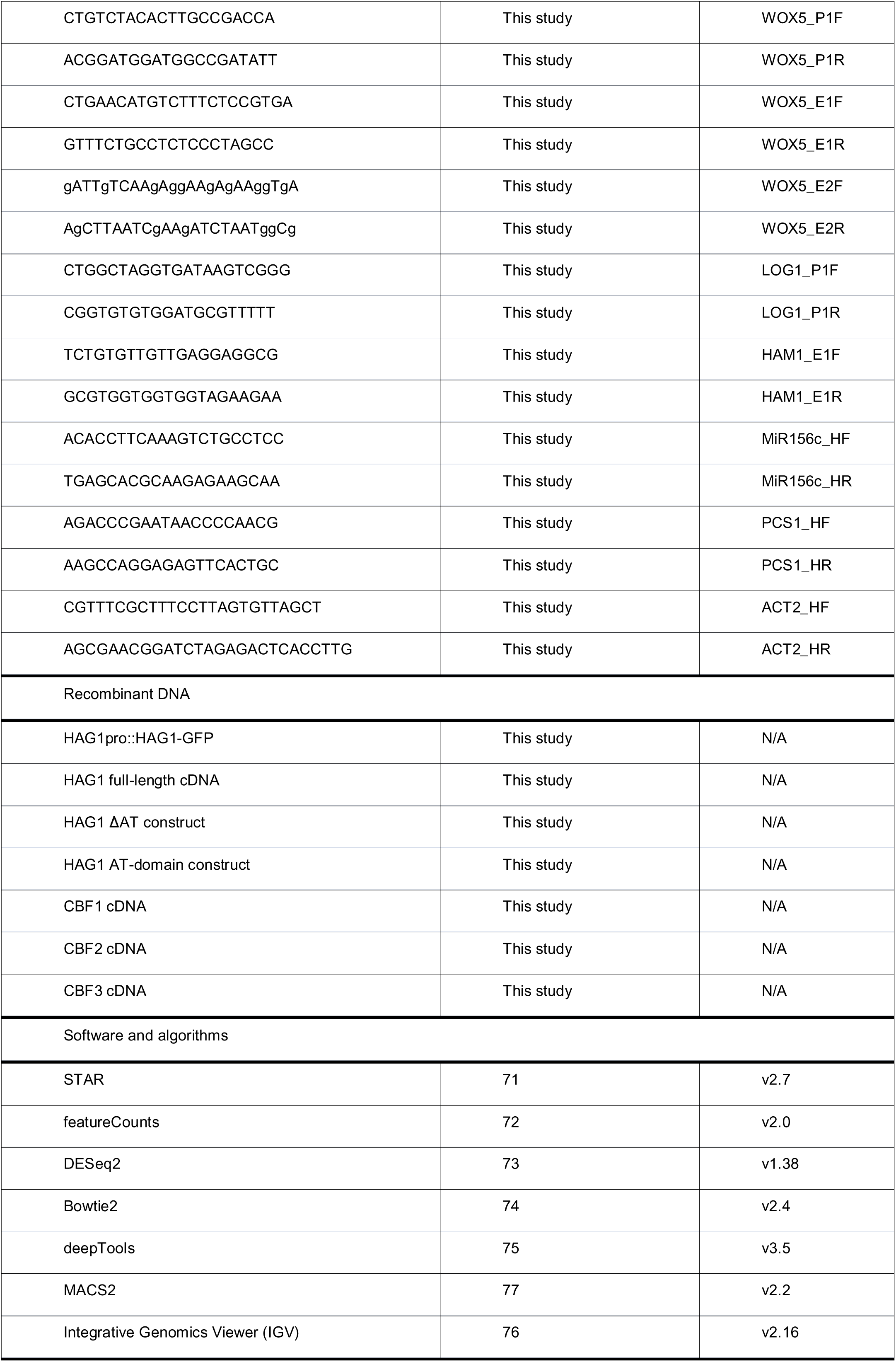
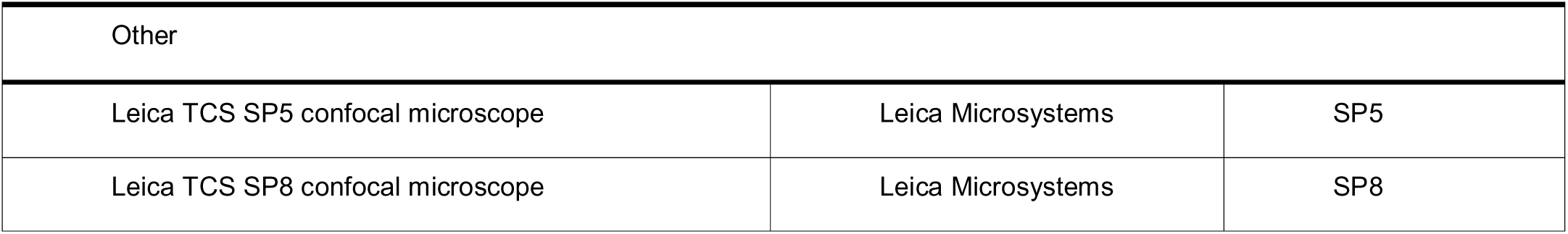

### EXPERIMENTAL MODEL AND STUDY PARTICIPANT DETAILS

#### Plant materials and growth conditions

Arabidopsis (*Arabidopsis thaliana*) plants were germinated on MS medium and grown at 22°C under short day (8 h light /16 h dark cycle) condition for 14 days. The *hag1-5* mutant (SALK_048427) is a T-DNA insertion mutant (62, 63). The *cbf1/2/3* triple mutants (*cbf1/2/3-97 and cbf1/2/3-93*) were Crisper-Cas9 generated lines (44). Other mutant or transgenic plants used in this study, i.e. *wox5-1* (64), *WOX5-pER8*, *WOX5pro:CRE/LOX-GUS* (26, 58), *WOX5pro::NLS-GFP* (53), *CBF3pro::NLS-mCherry, CBF3pro::CBF3-YFP* (55) and *FRI* Col (8) lines were reported previously. All mutant and transgenic plants used in this study were in the Col-0 background.

### METHOD DETAILS

#### Callus formation and regeneration assays

To induce callus, hypocotyls or first and second rosette leaves were cut from 14-day-old plants with microscissors (Natsume Seisakusho; MB-50-15) and their explants were incubated on phytohormone-free MS medium supplemented with 1% sucrose and 0.6% Gellan gum (Gelzan; Sigma-Aldrich) at 22℃ with continuous light. To induce shoot regeneration, 5-mm hypocotyl explants were cultured on SIM medium (65) at 22℃ with continuous light. Wound-induced callus phenotypes were recorded at 10 days after wounding and shoot regeneration phenotypes were recorded at 16 days after incubation on SIM. The projected area of callus was recorded by using Leica M165 C stereomicroscope.

#### Prolonged cold treatment

For cold exposure experiments, 14-day-old plants were transferred to 4℃ and grown under short day condition for varying durations prior to callus induction. Control samples were maintained under standard growth conditions. Following prolonged cold treatment, explants were immediately processed for regeneration assays or molecular analyses as described.

#### Histological analysis of resin-embedded semi-thin sections

Callus tissues were fixed overnight at 4°C in a solution containing 4% (w/v) paraformaldehyde and 2% (v/v) glutaraldehyde in a 0.05 M cacodylate buffer. After rinsing in the buffer, the specimens were post-fixed with 1% osmium tetroxide for two hours at room temperature, followed by dehydration in a graded ethanol series. The dehydrated samples were then infiltrated and embedded in EPON 812 epoxy resin (TAAB Laboratories Equipment Ltd.) and polymerized for 72 hours at 60 °C. Semi-thin sections (1 µm thick) were cut with a diamond histo knife (Diatome) on an ultramicrotome (EM UC7: Leicamicrosystems GmbH) and collected on glass slides. The sections were stained with a 0.05% toluidine blue solution (pH 7.0, Fujifilm Wako Co.) to enhance structural contrast and then rinsed with distilled water prior to imaging. The stained sections were imaged using an optical microscope (BX51M: Evident Scientific Co.) equipped with cellSens imaging software.

#### GUS staining and microscopy

*WOX5pro:CRE/LOX-GUS* lines (26, 58) were treated with 10 μM dexamethasone (DEX) for 14 days on SIM. GUS staining was conducted as previously described (66) with minor modifications. Samples were fixed in ice-cold 90% acetone for at least 30 min and rinsed twice with 50 mM sodium phosphate buffer (pH 7.2). Tissues were then incubated in staining solution containing 50 mM sodium phosphate buffer, 0.5 mM potassium ferricyanide, 0.5 mM potassium ferrocyanide, 0.1% Triton X-100, and 2 mM X-gluc. Vacuum infiltration was performed by applying vacuum until maximum pressure was reached, followed by rapid release; this cycle was repeated 3–4 times over a total of 10 min. Samples were then incubated at 37 °C overnight in the dark. Following staining, samples were cleared in 70% ethanol at room temperature until fully decolorized. Imaging was performed using a dissecting microscope.

#### Confocal microscopy

The GFP signals in *WOX5pro::NLS-GFP* were examined by using a TCS SP8 confocal microscope system with stander setting (Leica, https://www.leica-microsystems.com/products/confocal-microscopes), and the BiFC-YFP signals in agrobacterium transfected leaves were examined by using a TCS SP5 confocal spectral microscope imaging system.

To examine protein localization for higher resolution, the plants were cleared using ClearSeeAlpha and subsequently stained with SR2200. Fixation and clearing were performed according to a previously described protocol (67). For cell wall visualization, cleared samples were incubated in 0.1% (v/v) SR2200 (Renaissance Chemicals; the supplier’s stock solution was regarded as 100%) dissolved in ClearSeeAlpha for 24 hours. Images were acquired using a confocal laser scanning microscope (Zeiss LSM 880, Carl Zeiss, Germany) equipped with a C-Apochromat 40×/1.2 W Korr water-immersion objective. The objective zoom factor was 0.9×, and line averaging was set to 2 to improve the signal-to-noise ratio. Detector gain was adjusted to 400 (for SR2200), 750 (for GFP), and 700 (for mCherry). Laser power was 2% (405 nm), 15% (488 nm), and 4% (561 nm). The pinhole was set to 1 Airy unit. All imaging parameters were kept constant across samples. Z-stack images were acquired at 1 μm intervals and used to reconstruct optical cross sections using Fiji software.

#### Plasmid construction and plant transformation

The full-length coding sequences (CDS) of *HAG1/GCN5*, *CBF1*, *CBF2*, *CBF3*, and *CBF4* were PCR-amplified, cloned into the *pCR8/GW/TOPO* vector (Invitrogen) and subsequently recombined into the *pMDC43* vector to generate the *35Spro::GFP-CBF1* and *35Spro::GFP-CBF3* constructs. The *35Spro::HAG1-HA* constructs were generated by combining *HAG1-pCR8* and *pEAQ-HT* vector fused with *3xHA*. To generate the *HAG1pro::HAG1-GFP* and *HAG1pro::HAG1-3xFLAG* constructs, the *HAG1* genomic DNA sequence containing the ∼2 kb *HAG1* native promoter was PCR-amplified, cloned into the *pCR8/GW/TOPO* vector (Invitrogen), and then recombined into the *pGWB504* GFP vector or modified pEarlygate 301 vector with 3xFLAG sequence.

*HAG1pro::HAG1-GFP* transgenic plants were generated by transforming *HAG1pro::HAG1-GFP* into the *hag1-5* mutant by the floral dip method (68). To express *HAG1pro::HAG1-GFP* in the *cbf1/2/3* mutant background, *HAG1pro::HAG1-GFP* constructs were also introduced into the *cbf1/2/3-97* mutant by the floral dip method.

#### Bimolecular fluorescence complementation and co-immunoprecipitation assays

To generate the constructs for BiFC assays, full-length or truncated cDNA fragments of *HAG1/GCN5*, *CBF1*, *CBF2*, *CBF3*, and *CBF4* in the *pCR8/GW/TOPO* vector (Invitrogen) were recombined into the YN vector *pEarleyGate201-YN* or the YC vector *pEarleyGate202-YC*. The mCherry fused with nucleus localization signal (NLS) was also recombined into the *pEarleyGate201* vector, to indicate the nucleus. Constructed vectors were transiently transformed into *Nicotiana benthamiana* or the GVG-AvrPto *Arabidopsis* (69) leaves. Co-immunoprecipitation assays were performed as previously described (70). Anti-GFP (Santa Cruz Biotechnologies, catalog no. SC-9996; 1:3000 dilution) and anti-HA (Santa Cruz Biotechnologies, catalog no. SC-7392; 1:3000 dilution) antibodies were used as primary antibodies for Western blot, and the resulting signals were detected by using a Pierce ECL Western blotting kit (Pierce, https://www.lifetechnologies.com/).

#### RNA-seq analyses

For genome-wide expression analysis, RNA was isolated from control and prolonged cold treated seedlings of WT, *hag1-5* and *cbf1/2/3-97* plants using the QIAGEN RNeasy Plant Mini Kit (ID 74904) and treated with RNase-free DNase. RNA from at least three biological replicates was sequenced separately. Sequencing libraries were built using the KAPA RNA library preparation protocol. The libraries were sequenced on the Novoseq PE150 platform using a paired-end scheme (2 × 150 bp). Reads were mapped to the TAIR10 *Arabidopsis* genome using RNASTAR (71) with default settings. Normalized reads were calculated by featurecounts (72), the differential expressed genes and normalization ratio were analyzed by DEseq2 (73). The expression level was first normalized between WT, *hag1-5* and *cbf1/2/3-97*, and then normalized between control and prolonged cold treated samples among each background. The RNA-seq short read data have been submitted to the NCBI Gene Expression Omnibus (GEO) database (GSE297467). The log_2_ relative expression level and the normalized reads were also used to compare with the previous published data sets.

#### Chromatin immunoprecipitation assays

Chromatin extracts were prepared from seedlings treated with 1% formaldehyde. The chromatin was sheared to the mean length of 500 bp by sonication, and proteins and DNA fragments were immunoprecipitated using antibodies against anti-GFP (Abcam, catalog no. ab290). The DNA cross-linked to immunoprecipitated proteins was reversed, and then analyzed by real-time PCR using specific primers (Table S1). Percent input was calculated as follows: 2∧(Cq(IN)-Cq(IP))X100. Cq is the quantification cycle as calculated by the Biorad CFX Manager 3.1 based on the MIQE guidelines. Standard deviations represent at least 3 technical and 2 biological replicates. The variance in average data is represented by SEM (standard error of the mean). The SD (standard deviation), SEM determination and *p*-value were calculated using student’s paired *t*-test.

#### ChIP-seq and data analyses

Assays were performed as previously described (70). 2 ng of DNA from ChIP was used to ensure enough starting DNA for library construction. Two biological replicates were prepared and sequenced for each ChIP-seq experiment. The ChIP DNA was first tested by qRT-PCR and then used to prepare ChIP-seq libraries. End repair, adaptor ligation, and amplification were carried out using the KAPA DNA Library Prep kit (cat no. KR0961) according to the manufacturer’s protocol. The Novoseq PE150 was used for high-throughput sequencing of the ChIP-seq libraries. The raw sequence data were processed using the GAPipeline Illumina sequence data analysis pipeline. Bowtie2 (74) was then employed to map the reads to the *Arabidopsis* genome (TAIR10). The mapped reads were retained for further analysis. The alignments were first converted to Wiggle (WIG) files using deepTools (75). The data were then imported into the Integrated Genome Viewer (IGV) (76) for visualization. The H3Ac peaks were analyzed by MACS2 (77). The ChIP-seq short read data have been submitted to the NCBI Gene Expression Omnibus (GEO) database (GSE297486).

### QUANTIFICATION AND STATISTICAL ANALYSIS

All quantitative data are presented as mean ± standard error of the mean (SEM) or boxplots. Statistical analyses were performed using standard statistical software. The exact number of biological replicates (n), definition of n, and statistical tests used for each experiment are specified in the corresponding figure legends.

For comparisons between two groups, student’s *t*-tests were applied. For multiple group comparisons, two-way ANOVA followed by appropriate post hoc tests was used. No data were excluded unless pre-established technical failure criteria were met. Differences were considered statistically significant at *p* < 0.05, unless otherwise indicated.

### ADDITIONAL RESOURCES

No additional external resources were generated for this study.

