## Supplemental Figures for "Prolonged cold exposure enhances regeneration potential in Arabidopsis"

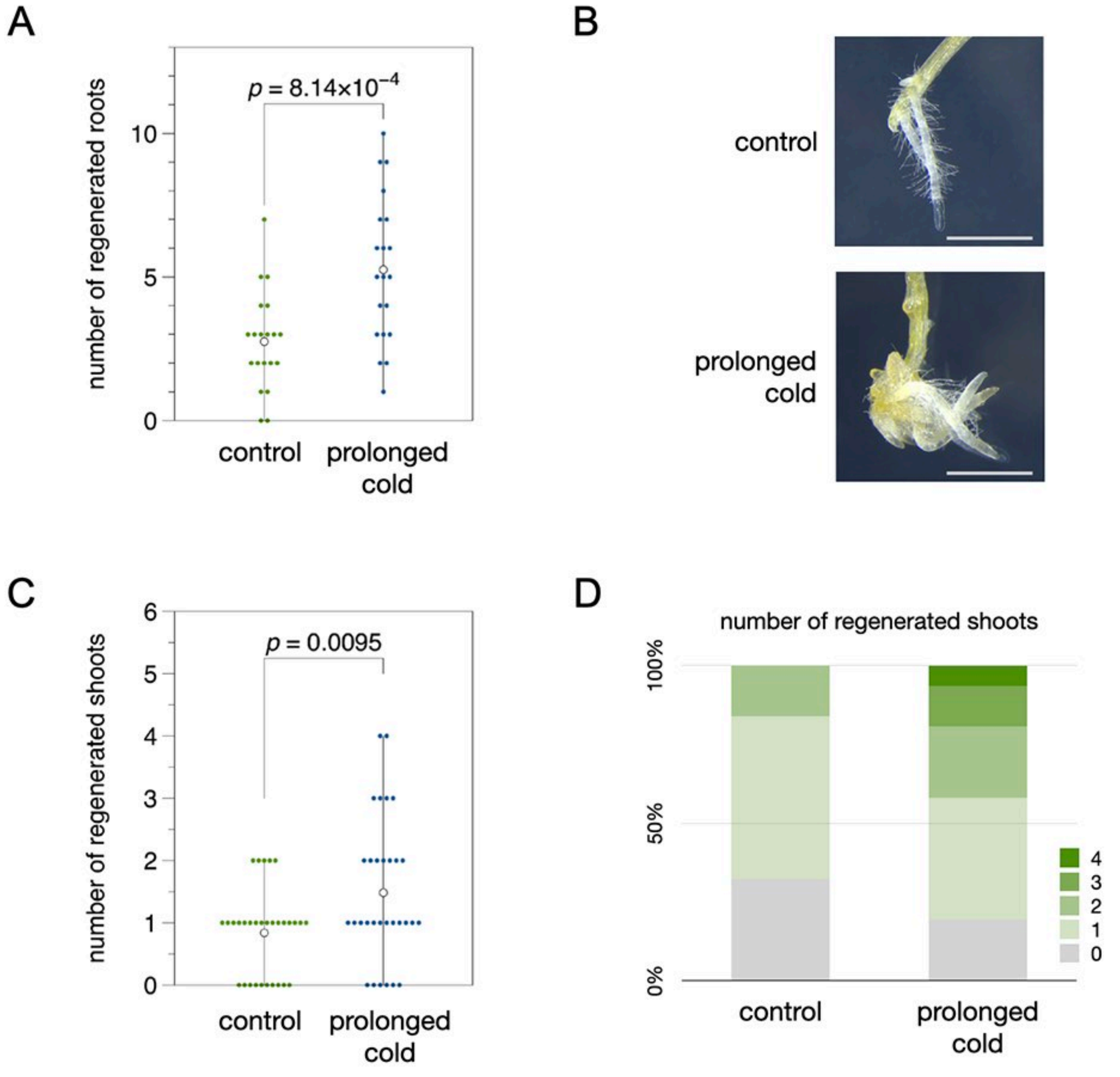


**Figure S1. Prolonged cold treatment promotes root and shoot regeneration in Arabidopsis. Related to Figure 1.**
(A, B) Quantitative data (A) and phenotypes (B) of regenerated adventitious roots from hypocotyl after wounding. Scale bars, 1 mm. Control or prolonged cold treated samples were cultured for 5 days after cutting. (C, D) The quantitative data (C) and distribution ratio (D) of regenerated shoots from control or prolonged cold treated hypocotyl explants cultured in shoot inducting medium (SIM). The number of regenerated shoots were calculated at 16 days after culture. All experiments were performed at least three times with comparable results.

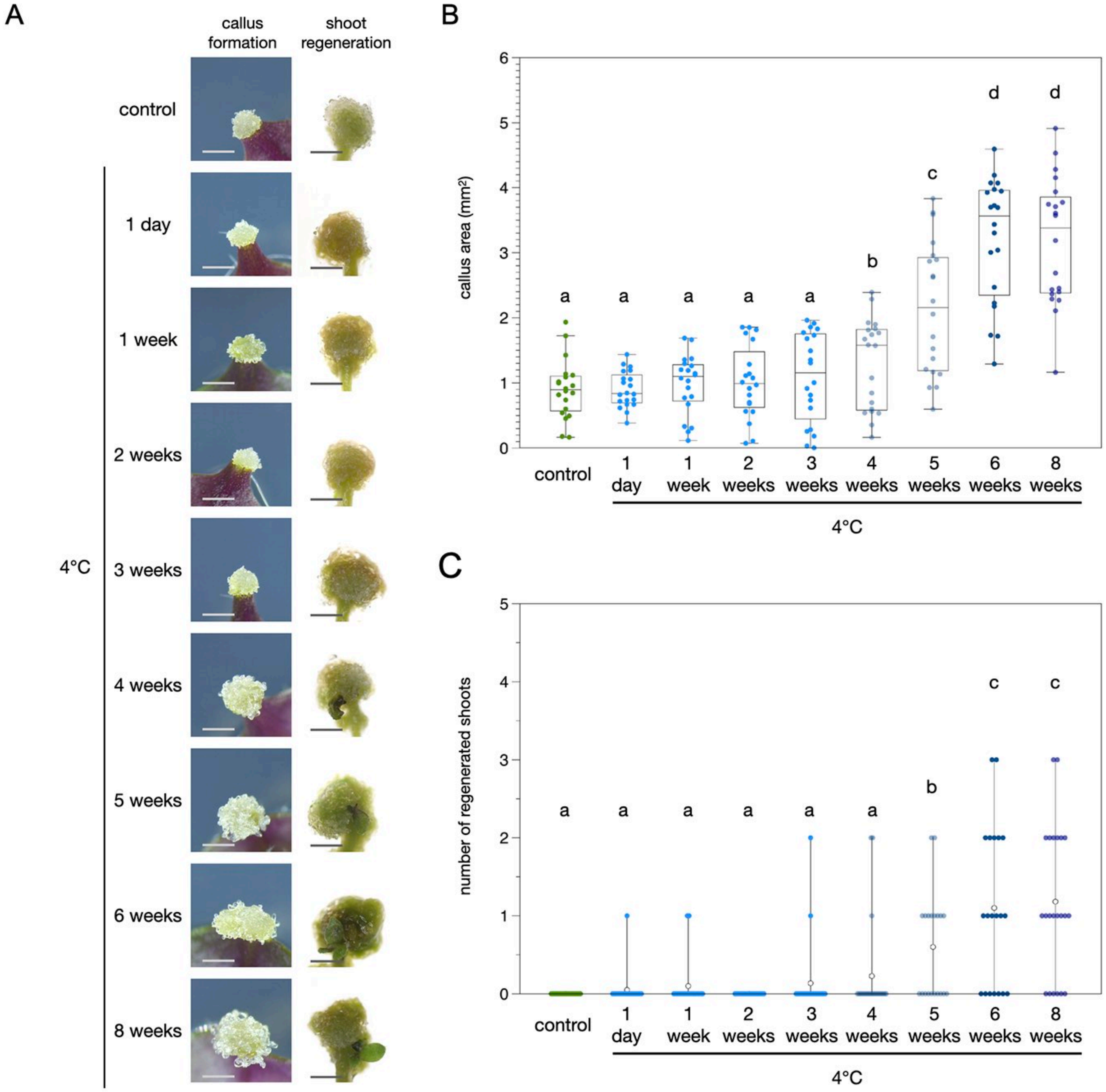


**Figure S2. Long-term cold treatment is required to promote callus formation and shoot regeneration in Arabidopsis. Related to Figure 1.**
(A) Phenotypes of wound-induced callus formation and shoot regeneration in control seedlings and seedlings exposed to cold treatment for different durations. (B, C) Quantitative data of wounding induced callus formation (B) and shoot regeneration (C) in control and cold-treated seedlings. Letters indicated statistical significance based on a two-factor ANOVA with Tukey's HSD post hoc analysis ($p < 0.05$). All experiments were performed at least three times with comparable results. Scale bars, 1 mm.

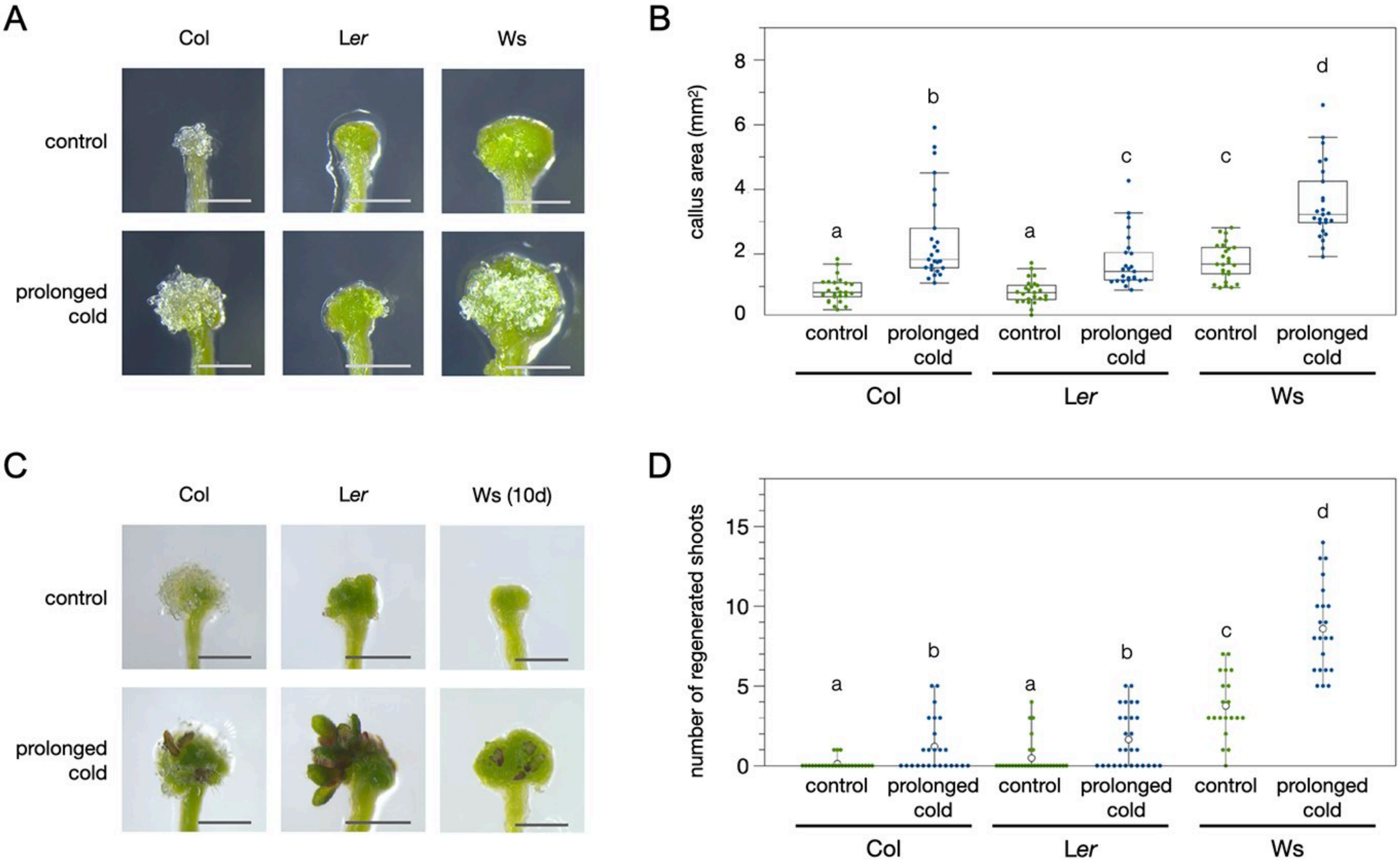


**Figure S3. The prolonged cold promoted regeneration is consistently observed in different ecotypes of Arabidopsis. Related to Figure 1.**
(A) Light microscope images of callus forming from wounded hypocotyls. Scale bars, 1 mm (B) Quantitative data of callus area. Samples sharing the same letters did not show statistically significant differences. (C) Light microscope images of regenerating shoots from hypocotyl explants cultured on shoot inducting medium (SIM). The images of Col and L*er* were collected on 16 days after culture, and the images of Ws were collected on 10 days after culture. Scale bars, 1 mm. (D) Quantitative data of shoot regeneration. The number of regenerating shoots was quantified on 16 days after culture. Letters indicated statistical significance based on a two-factor ANOVA with Tukey's HSD post hoc analysis ($p < 0.05$). All experiments were performed at least three times with comparable results.

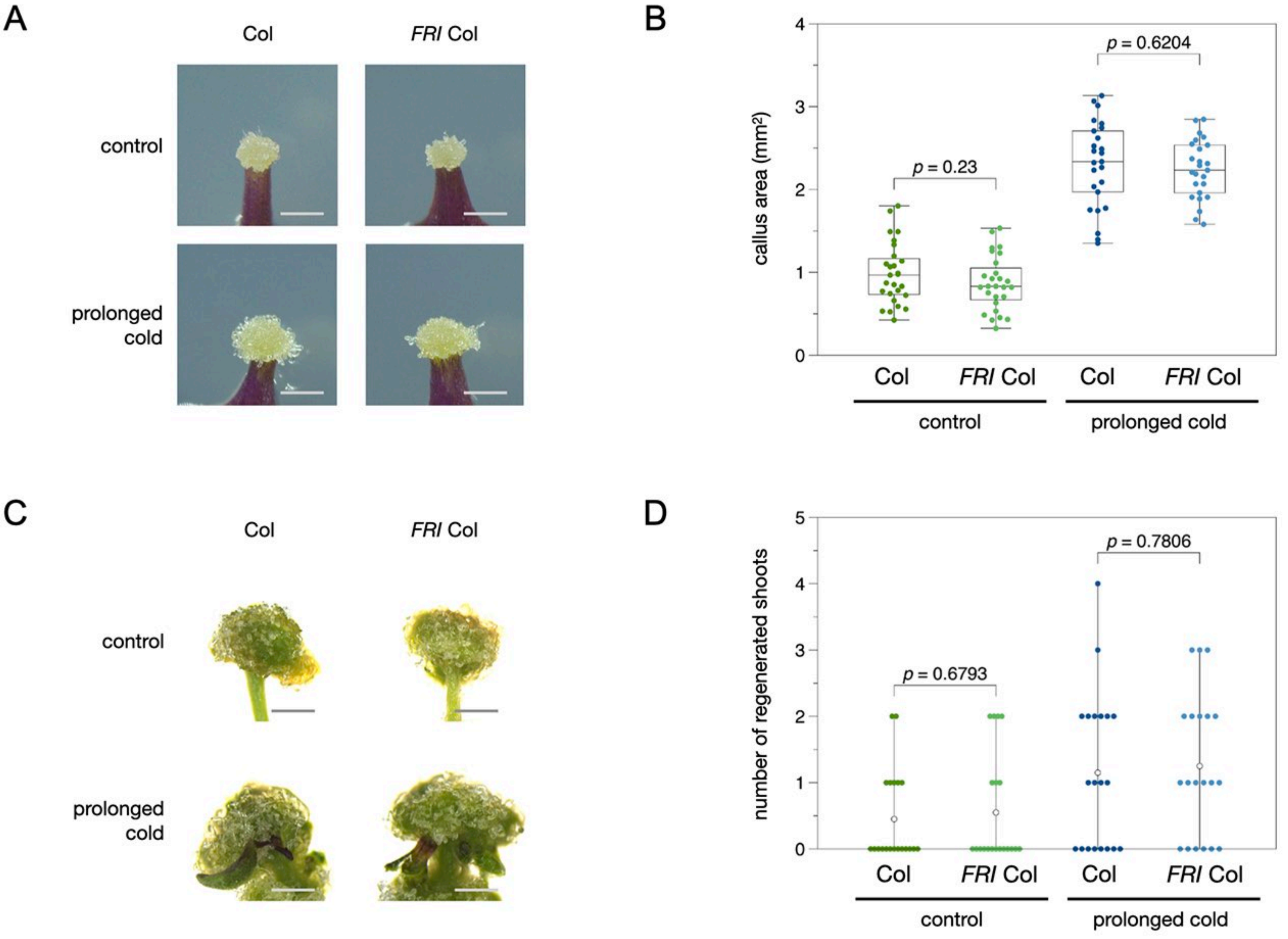


**Figure S4. The prolonged cold promoted regeneration is not affected by FRI. Related to Figure 2.**
(A-D) Phenotypes and Quantitative data of wounding induced callus (A, B) and shoot regeneration (C, D) in WT-col and FRI Col. All experiments were performed at least three times with comparable results. Scale bars, 1 mm.

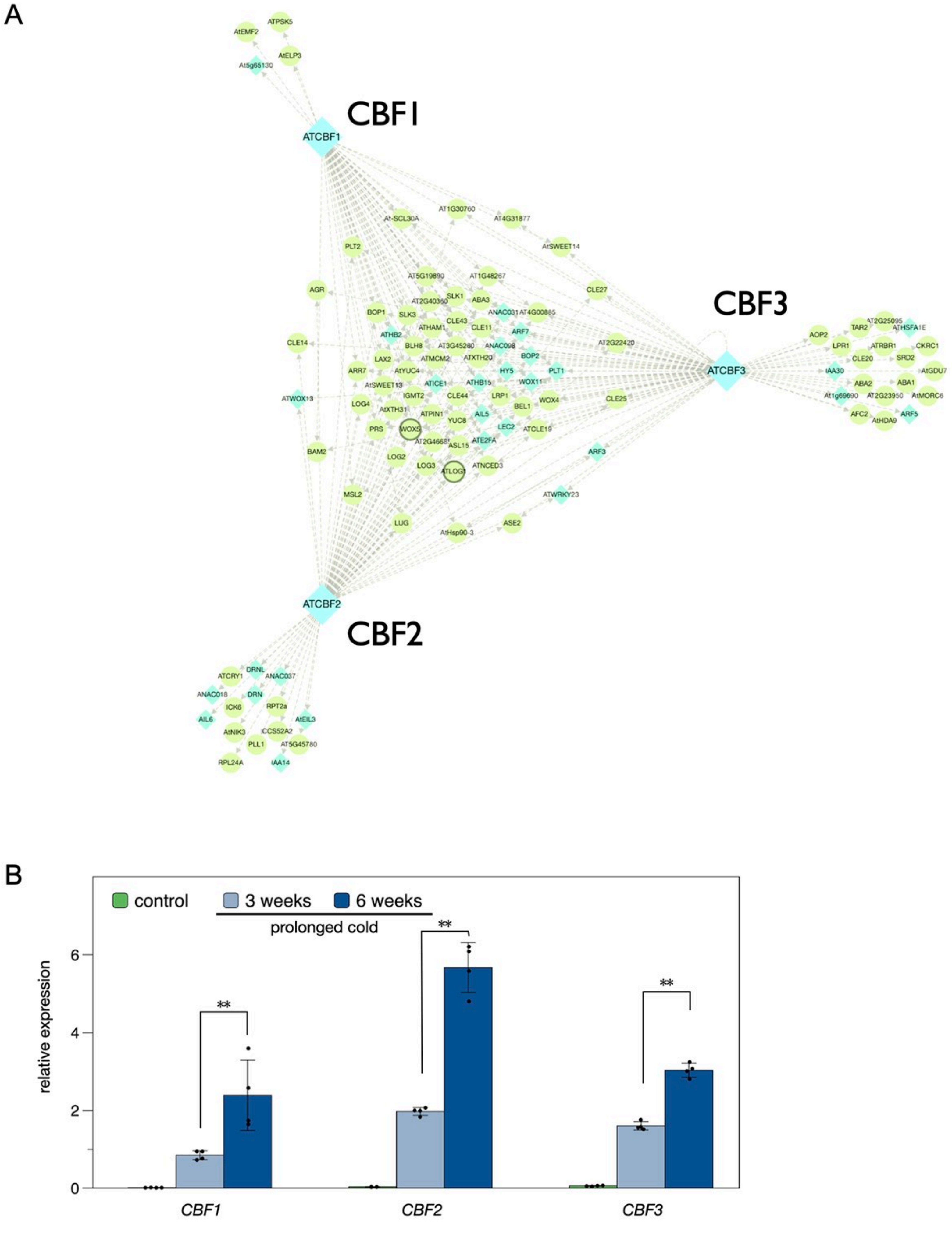


**Figure S5. CBF1/2/3 are putative upstream regulators of the prolonged cold activated expression of regeneration genes. Related to Figure 2.**
(A) The genes up-regulated by prolonged cold among the regeneration associated genes (Data. S2) were sent to the gene regulatory network (GRN) analysis tool TF2Network (35) to predict putative upstream regulators of the prolonged cold promoted regeneration. (B) Expression of *CBF1*, *CBF2* and *CBF3* in control and seedlings treated with 3 weeks or 6 weeks of prolonged cold. The expression level was analyzed by qRT-PCR, using *PP2A* as an internal control. Error bars indicate standard derivations. At least three independent biological replicates were used for the analysis. ** $p < 0.005$ (student's t-test)

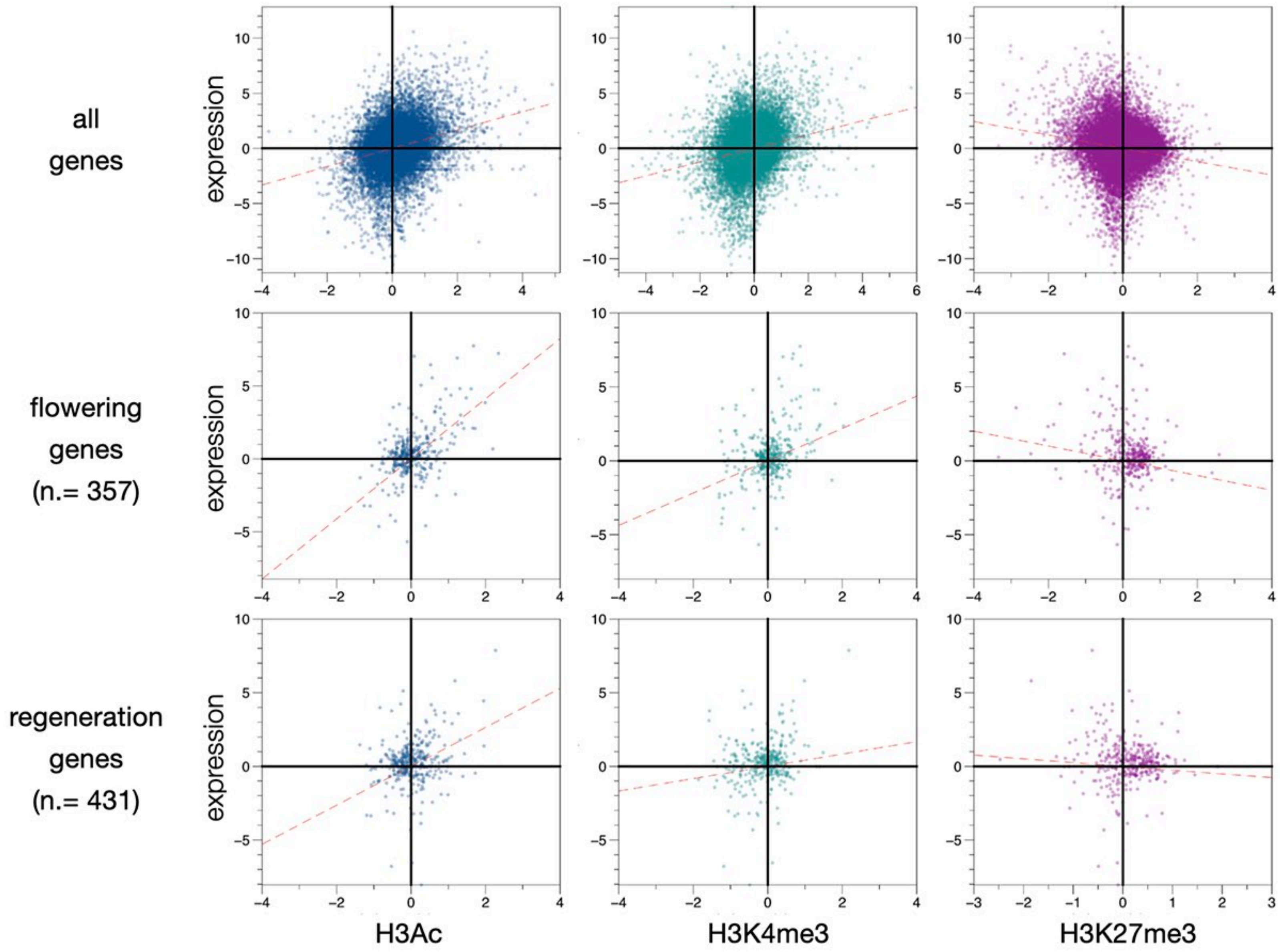


**Figure S6. The general expression changes of the regeneration associated genes after prolonged cold are correlated with changes in H3Ac change. Related to Figure 2.**
The X-Y scatter plots and ablines showing the correlation between expression level changes with H3Ac, H3K4me3 or H3K27me3. The ablines were calculated by the genes that showed significant changes in histone modification levels ($\log_2$ value <-1 or >1). These data show that among all Arabidopsis coding genes, gene expression after prolonged cold is positively correlated with H3Ac, H3K4me3, and negativity correlated with H3K27me3. Similar correlations were also found in 357 flowering associated genes (Table S5) which was used as a control for smaller total gene number. However, the expression level change of 431 regeneration associated genes (Table S1) was most strongly correlated with H3Ac level changes compared to H3K4me3 or H3K27me3.

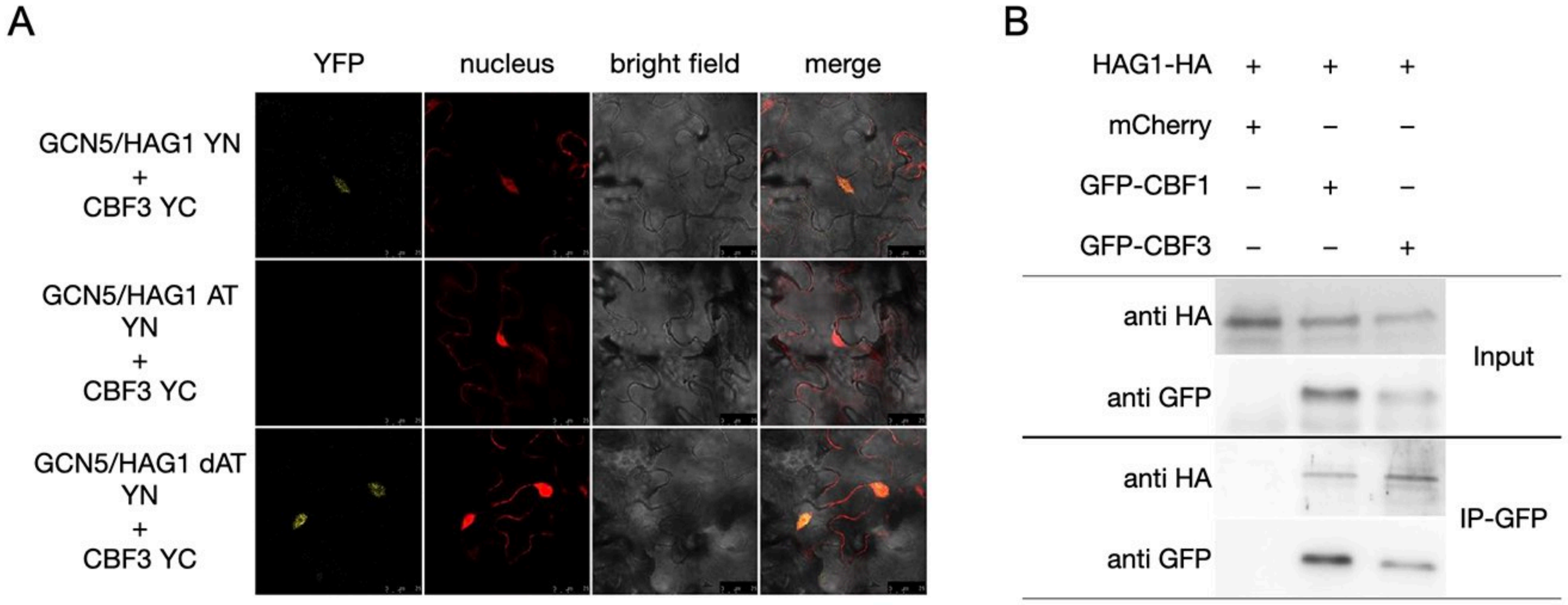


**Figure S7. HAG1 and CBFs interact in Arabidopsis cells. Related to Figure 2.**
(A) BiFC assays in Arabidopsis leaves showing the interaction of HAG1 with CBF3 in living cells. HAG1 and the CBF proteins were fused with the N terminus (YN) or C terminus (YC) of YFP and co-delivered into Arabidopsis leaves by agrobacterium infiltration. The nucleus was indicated by mCherry carrying a nuclear localization signal. Different deletion form of HAG1 were used to examine the interaction with CBF3. Scale bars, 10 μm. (B) Co-IP of HAG1-HA with CBF1-GFP and CBF3-GFP co-delivered into Arabidopsis leaves by agrobacterium infiltration. Western blot experiments were performed with the indicated antibodies.

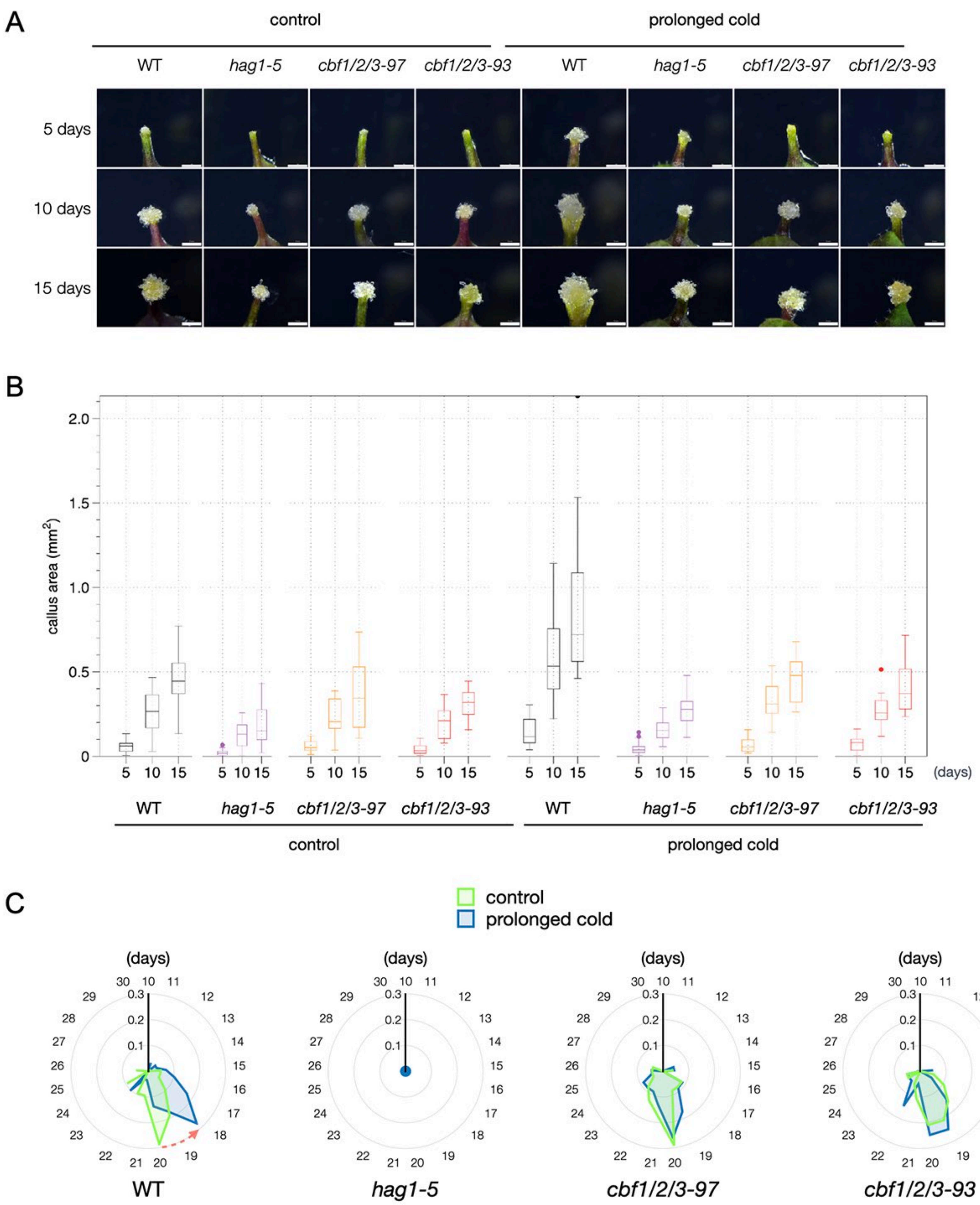


**Figure S8. The *hag1* and *cbf* mutants display defects in prolonged cold promoted callus formation. Related to Figure 3.**

(A, B) Phenotypes (A) and quantitative data (B) of wounding induced callus in control or prolonged cold treated WT, *hag1* and *cbf1/2/3* mutant plants. Scale bars, 1 mm. (C) Quantitative data of shoot regeneration in control or prolonged cold treated hypocotyl explants from WT, *hag1-5* and *cbf1/2/3-97* and *cbf1/2/3-93* mutant plants cultured on shoot-inducting medium (SIM). Shoot regeneration was scored up to 30 days on SIM and shown as a ratio of increased shoots per day divided to total explant number. All data were performed with at least three independent experiments with comparable results.

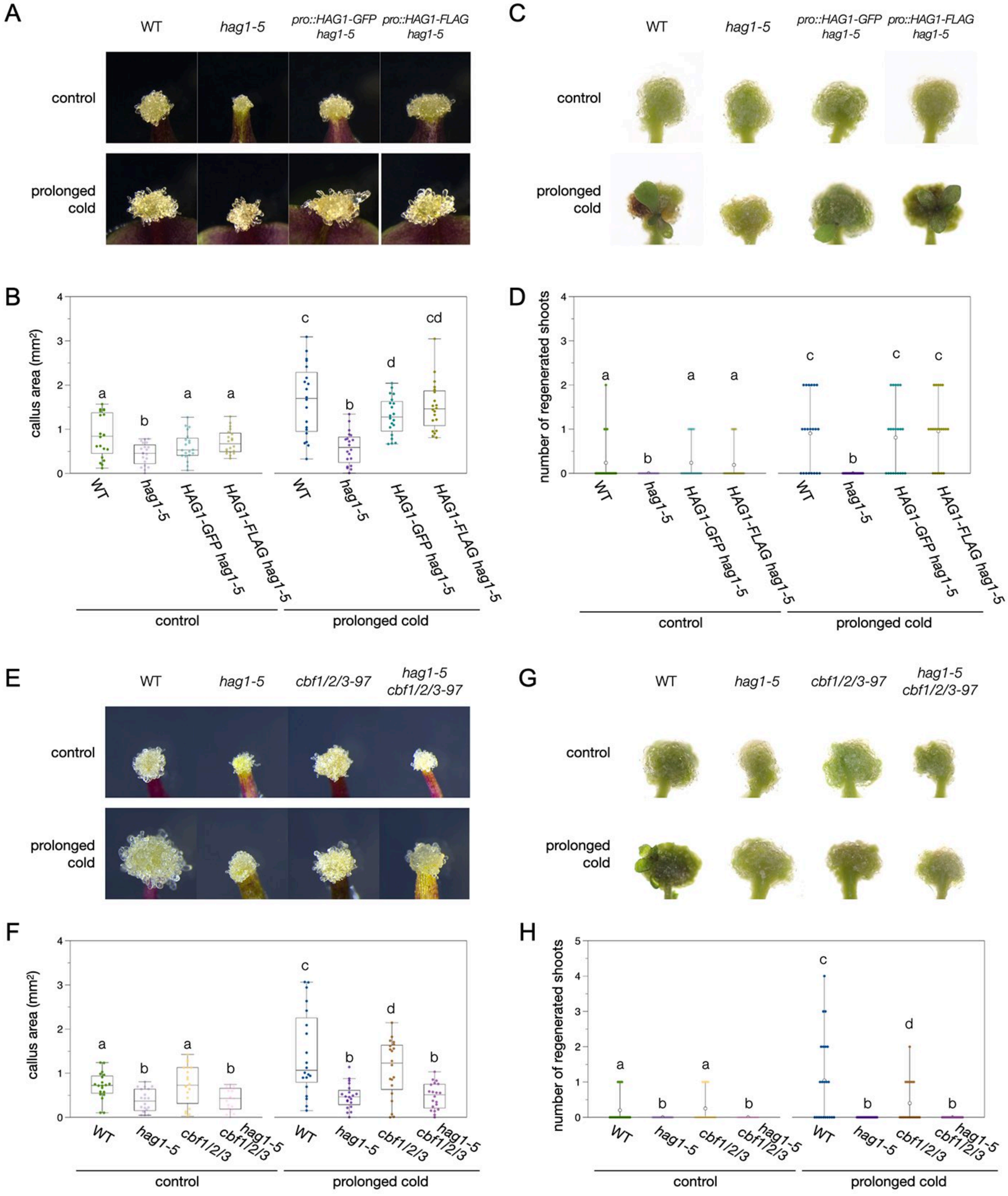


**Figure S9. The CBFs-HAG1 regulatory module is involved in the regulation of prolonged cold promoted regeneration. Related to Figure 3.**
(A-D) Phenotypes and Quantitative data of wounding induced callus (A, B) and shoot regeneration (C, D) in control or prolonged cold treated WT, *hag1-5* and *pro::HAG1-GFP hag1-5 and pro::HAG1-FLAG hag1-5* lines. (E-H) Phenotypes and Quantitative data of wounding induced callus (E, F) and shoot regeneration (G, H) in control or prolonged cold treated WT, *hag1-5*, *cbf1/2/3* and *hag1-5 cbf1/2/3* mutant plants. Letters indicated statistical significance based on a two-factor ANOVA with Tukey's HSD post hoc analysis ($p < 0.05$). All experiments were performed at least three times with comparable results. Scale bars, 1 mm.

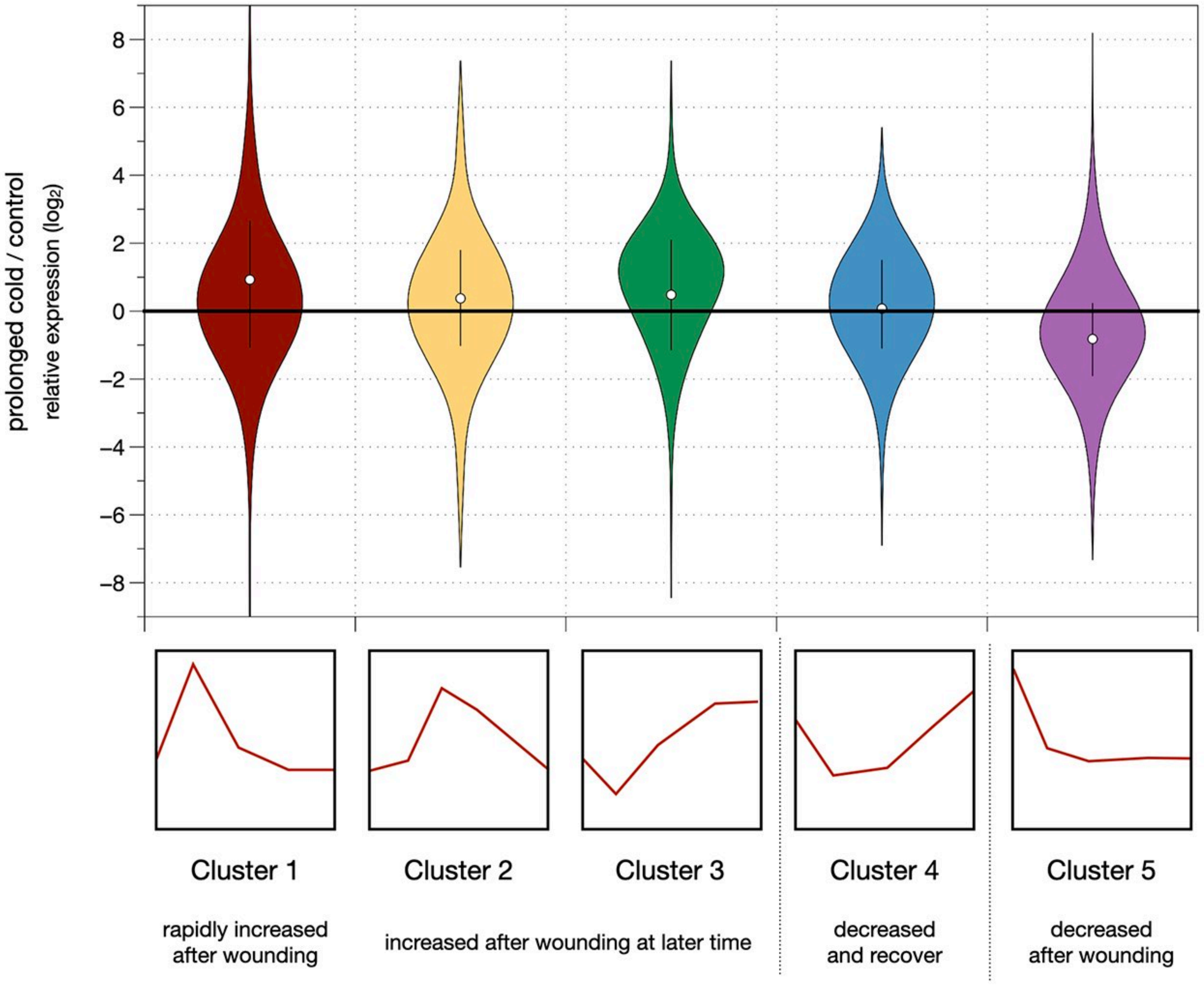


**Figure S10. Transcriptional changes after the prolonged cold treatment are positively correlated with those after wounding. Related to Figure 3.**
The violin plots showing the relative expression after prolonged cold treatment among different types of wounding response genes. The clusters were classified following previous study (20). The genes with $p < 0.0001$ were used for analysis (Data S1). White dots indicated average value and the straight line indicated 50 percentile value. The results showing the general relative repression pattern after prolonged cold treatment is increased in wound-induced genes and decreased in wound-reduced genes.

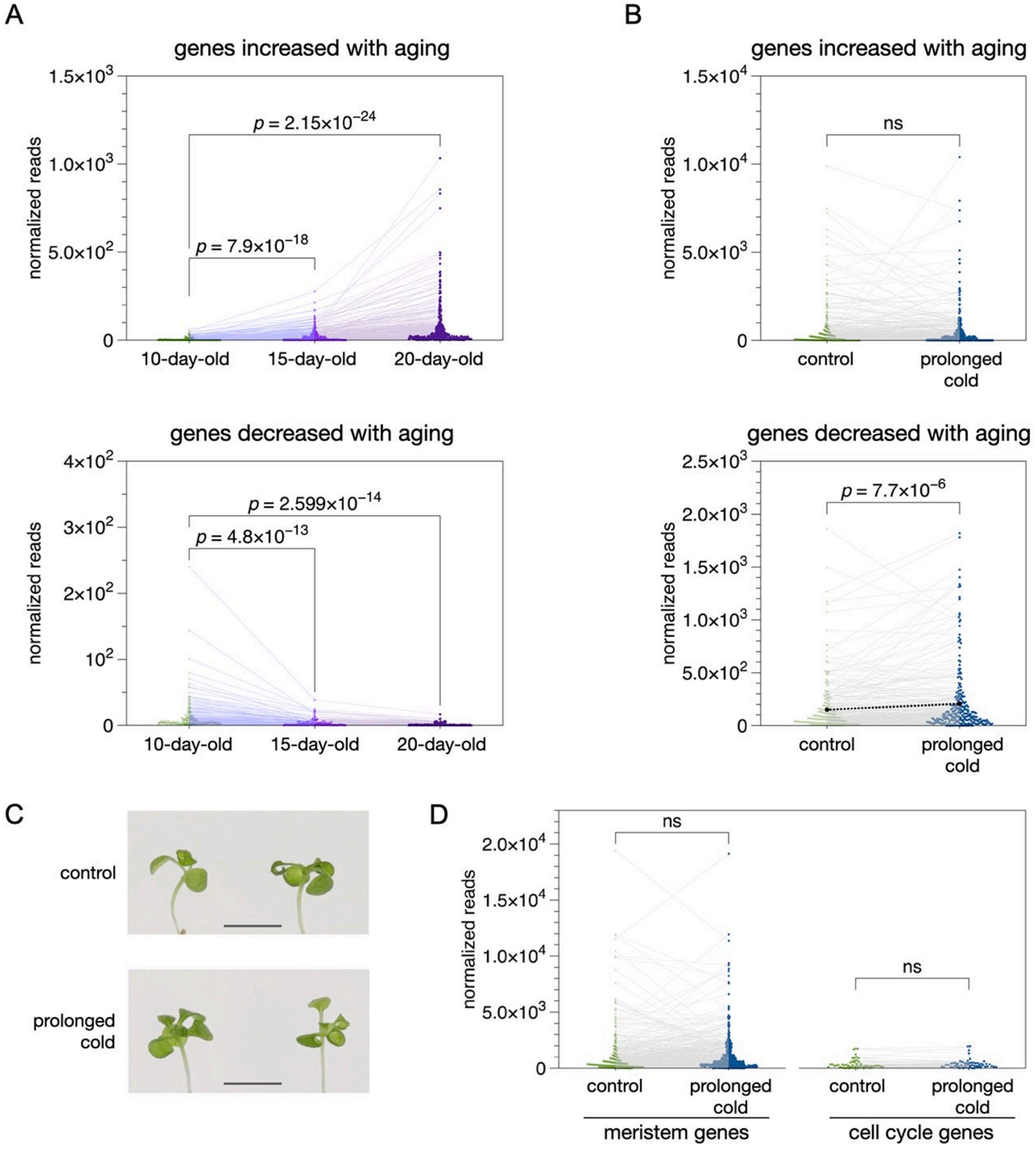


**Figure S11. The expression pattern changes of aging-induced or decreased genes are not associated with prolonged cold treatment. Related to Figure 4.**
(A) The normalized reads showing the expression levels of the genes increased or decreased with aging in Arabidopsis. Each dot indicated the expression level of aging -induced or -decreased genes in 10-, 15- and 20-day-old plants based on a published study (45). (B) The normalized reads showing the expression levels of aging -induced or -decreased genes after prolonged cold. The black dots and the dotted line showed the average level of the genes decreased with aging in control or prolonged cold treated plants. (C) The Morphological analysis in aerial tissues in the prolonged cold-treated and control plants. Scale bars, 10 mm. (D) The normalized reads showing that the overall expression pattern of genes associated with meristem development or cell cycle regulation was not significantly altered after the prolonged cold. ns = no significant difference ($p > 0.05$).

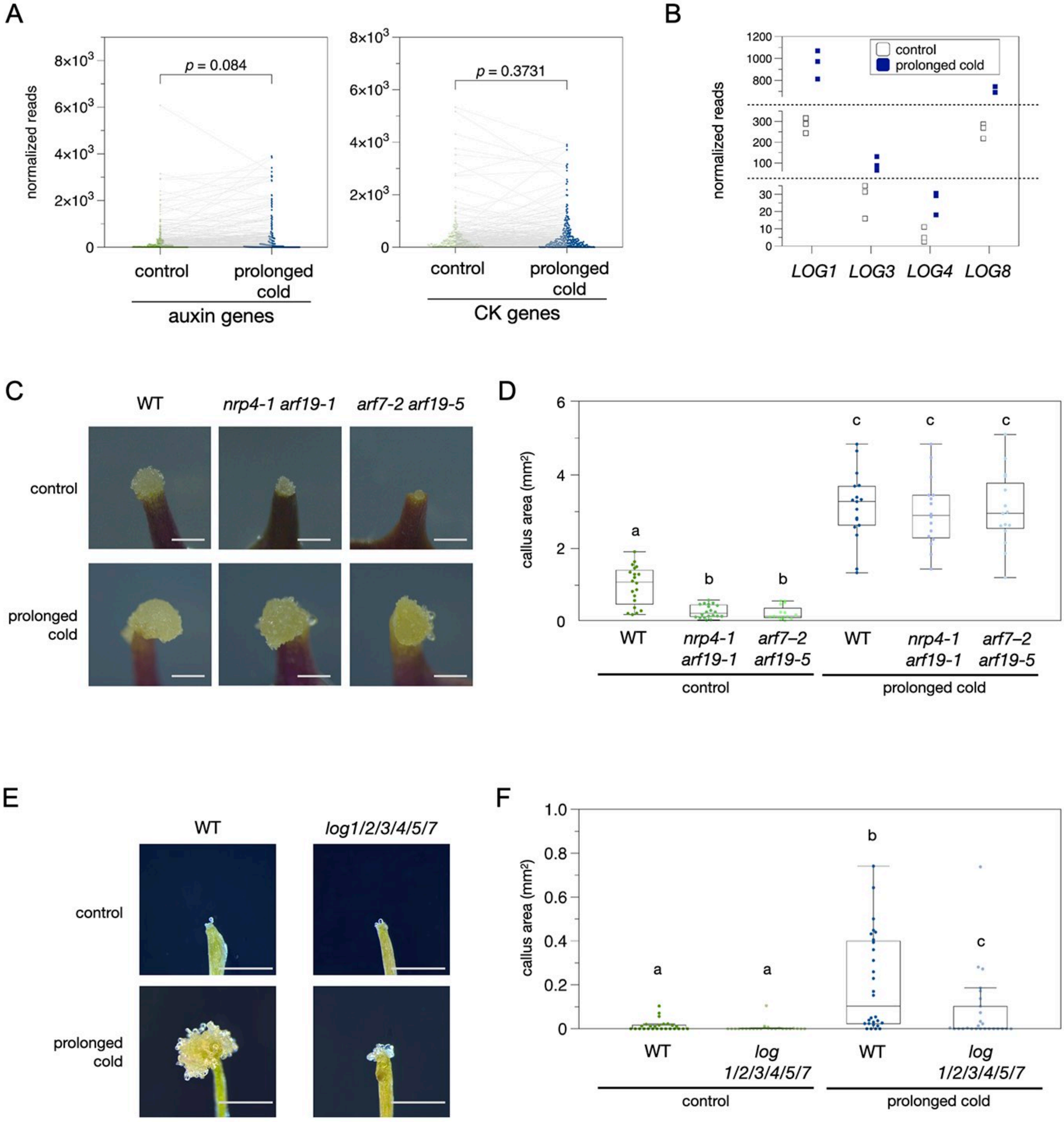


**Figure S12. Prolonged cold-promoted regeneration is associated with cytokinin-mediated pathway. Related to Figure 5.**

(A) The normalized reads showing that the overall expression pattern of genes associated with auxin and cytokinin (CK) was not significantly altered after the prolonged cold. (B) The normalized reads of *LOG1*, *LOG3*, *LOG4*, *LOG8* in the control and prolonged cold treated plants. (C, D) Phenotypes (C) and quantitative data (D) of the callus induced by wounding in WT, *nrp4-1 arf19-1*, and *arf7-2 arf19-5* mutant lines. (E, F) Phenotypes (E) and quantitative data (F) of shoot regeneration in WT and *log1/2/3/4/5/7* mutant lines. Letters indicated statistical significance based on a two-factor ANOVA with Tukey's HSD post hoc analysis ($p < 0.05$). All experiments were performed at least three times with comparable results. Scale bars, 1 mm.

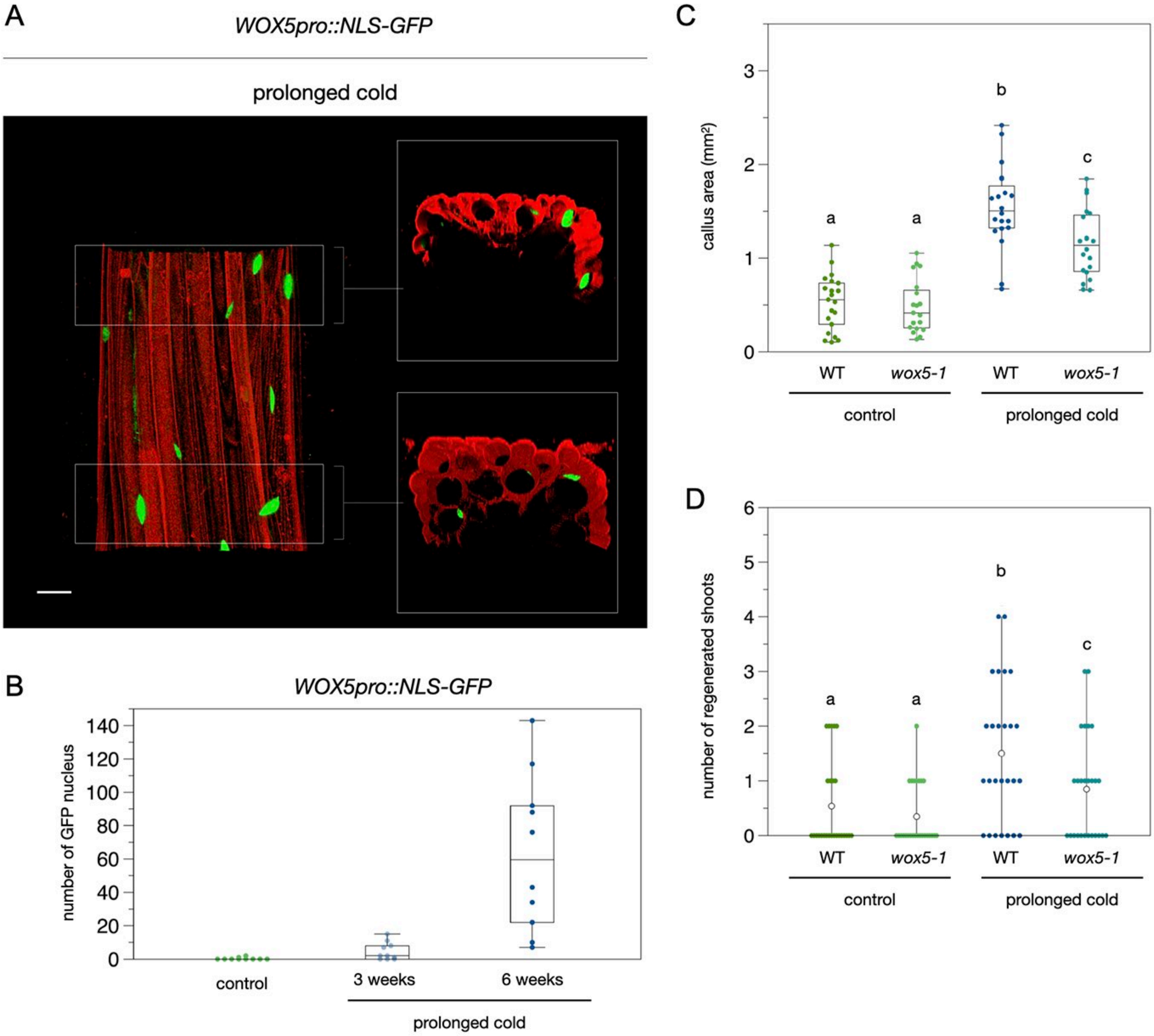


**Figure S13. Increased *WOX5* expression during prolong cold contributes to callus formation and shoot regeneration. Related to Figure 6.**

(A) The 3D reconstruction of confocal microscope images visualizing the NLS-GFP expression driven by the *WOX5* promoter in the Arabidopsis hypocotyl treated with prolonged cold. The GFP signal was marked in green and PI staining was marked in red. Scale bars,100 μm. (B) The number of the GFP positive nucleus in the *WOX5pro::NLS-GFP* plants in control or after prolonged cold treatment. The GFP positive nucleus were counted within 1 cm of hypocotyls. (C, D) Quantitative data of wounding induced callus (C) and regenerated shoots (D) in control or prolonged cold treated WT and *wox5-1* mutant plants. Letters indicated statistical significance based on a two-factor ANOVA with Tukey's HSD post hoc analysis ($p < 0.05$). All experiments were performed at least three times with comparable results.
